# Astrocytic chordin-like 1 (Chrdl1) is re-engaged early after ischemic stroke to regulate region-dependent GluA2 levels and neuronal vulnerability

**DOI:** 10.64898/2026.08.18.745607

**Authors:** BR Boyle, RB Hastings, A Patel, AJ Gleichman, ST Carmichael, E Blanco-Suarez

## Abstract

Neuronal vulnerability to ischemic stroke varies markedly across brain regions, yet the mechanisms underlying this selective susceptibility remain poorly understood. Here, we show that the developmental astrocytic protein Chordin-like 1 (Chrdl1) is repurposed after ischemic injury to regulate neuronal vulnerability. Chrdl1 expression stabilizes GluA2-containing AMPA receptors, limits delayed apoptotic neuronal death, and preserves hippocampal function early after focal ischemic stroke, whereas sustained Chrdl1 expression does not improve long-term recovery. These findings identify an unexpected neuroprotective role for Chrdl1 during acute ischemia that contrasts with its previously described function as a limiter of synaptic plasticity during recovery. Our work reveals that developmental astrocyte-derived signaling can be redeployed after brain injury, with distinct functions depending on the stage of stroke and region-dependent endogenous expression that together determine whether a conserved neuroprotective mechanism is engaged after ischemic stroke.

## INTRODUCTION

Recovery from brain ischemic injury requires a delicate balance between preserving vulnerable neurons immediately after the insult while allowing circuit remodeling that restores function. Some molecular pathways that are typically active during development may be reactivated in response to injury^1^, yet whether these reactivated developmental mechanisms promote repair or impede recovery likely depends on when and where they are engaged. How developmental mechanisms are repurposed following ischemic injury remains a major unanswered question, with important implications for neuronal vulnerability and functional recovery after stroke.

Astrocytes regulate synaptic maturation, receptor composition, and plasticity through secreted proteins during development^2^, and are increasingly recognized as important players in stroke pathophysiology, beyond border formation and neuroinflammation^3,4^. Following ischemic injury, astrocyte signals present time-dependent functions, influencing diverse biological processes throughout the different stages of stroke progression^4^, and can either promote recovery or contribute to maladaptive remodeling that compromises neuronal survival and functional restoration^4,5^, especially on more vulnerable neuronal populations.

One key determinant of neuronal vulnerability to ischemic injury is the GluA2 subunit of the AMPA receptors (AMPARs), which renders the AMPAR impermeable to Ca^2+^. Shortly after ischemic injury, GluA2-containing AMPARs are rapidly removed from the neuronal surface in particularly vulnerable populations, such as hippocampal neurons^6,7^. This alters the ratio of GluA2-containing versus GluA2-lacking AMPARs, promoting excessive Ca^2+^ influx through GluA2-lacking, Ca^2+^-permeable AMPARs and exacerbating excitotoxicity^8,9^. This results in intracellular signaling cascades that activate caspase-3 and other molecules that ultimately lead to delayed apoptotic cell death^10^. When these pathways are inhibited, or GluA2 levels are unaffected after ischemic insult, neuronal survival is significantly increased^6,10–12^.

While the neuronal mechanisms underlying the rapid stroke-induced internalization of GluA2 in hippocampal neurons have been previously characterized^6,7,13^, considerably less is known about the non cell-autonomous mechanisms that regulate this process. Astrocytes, as previously mentioned, regulate AMPAR trafficking to promote synaptic formation, maturation and stabilization during development. One such astrocyte-secreted protein is Chordin-like 1 (Chrdl1) which promotes synaptic maturation by increasing synaptic GluA2-containing AMPARs and limiting synaptic plasticity in the mouse cortex^14^. We have previously shown that Chrdl1 is robustly upregulated in peri-infarct astrocytes following ischemic stroke in cortical regions^12,15^, and that genetic deletion of Chrdl1 improves functional recovery only during the subacute and chronic phases after stroke, whereas no differences in motor performance were observed during the acute phase^12^.

These findings raised an unexpected possibility. Although Chrdl1 limits synaptic plasticity during stroke recovery, it may instead serve as an early astrocyte-derived neuroprotective signal by stabilizing GluA2- containing AMPARs before excitotoxic signaling is initiated, with the potential to later constrain the plasticity required for synaptic repair. More broadly, whether developmental astrocyte-derived mechanisms are repurposed during ischemic injury to regulate neuronal vulnerability in a spatiotemporal manner remains unknown.

Here, we show that at early post-stroke stages, astrocytic Chrdl1 expression stabilizes GluA2-containing AMPARs, limits delayed apoptotic neuronal death and preserves hippocampal function following focal hippocampal ischemic stroke. In contrast, sustained Chrdl1 expression does not improve long-term recovery. Together, our findings reveal that developmental astrocyte-derived signals may be repurposed after ischemic injury, with distinct functions depending on the stage of stroke and the affected brain region.

## RESULTS

### Selective neuronal vulnerability after ischemic stroke is associated with differential GluA2 levels

Previous work demonstrated that cortical and hippocampal neurons differentially regulate AMPAR subunit composition following oxygen and glucose deprivation (OGD), with rapid loss of GluA2 occurring selectively in hippocampal neurons^6,9^, suggesting region-specific mechanisms of AMPAR remodeling. To determine whether these region-specific differences in GluA2 levels are preserved *in vivo*, we employed a focal ischemic stroke model using photothrombosis to induce localized ischemic injuries in discrete brain regions^12,16,17^. Focal ischemic strokes were generated in the motor cortex or hippocampus of adult male mice and the levels of GluA2 in the peri-infarct regions were analyzed 2 hours post-surgery (hps, sham or stroke). Successful stroke induction and anatomical localization were confirmed immediately prior to tissue collection using MRI (Figure 1A, S1A).

**Figure 1.**
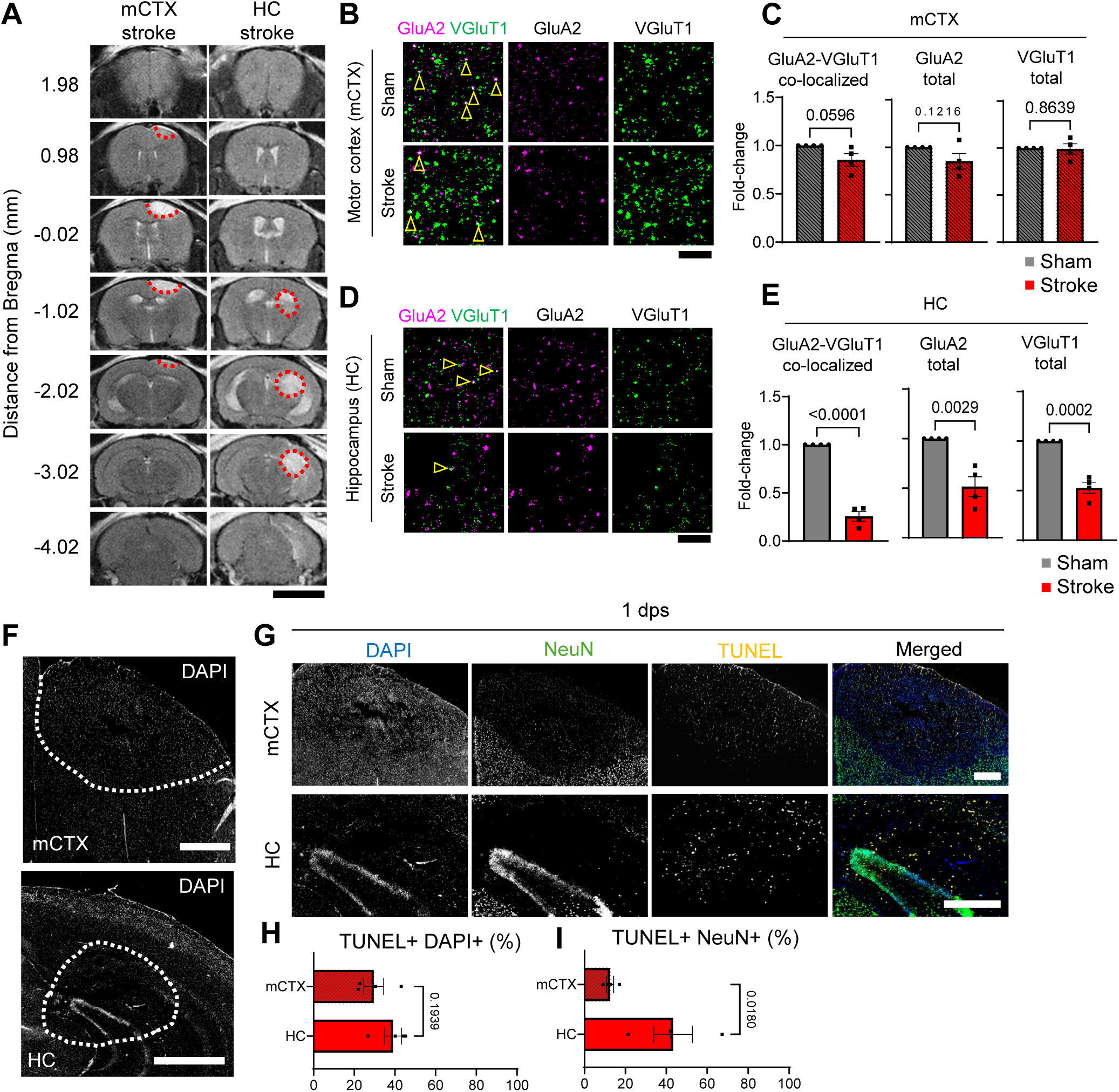
***Selective neuronal vulnerability after ischemic stroke is associated with differential GluA2 levels.*** A) Representative images of coronal T2-weighted MRI scans spanning the whole injury in the mCTX or hippocampus (from 1.98 mm rostral to -4.02 mm caudal to bregma) of male WT animals at 2 hours post-surgery (hps). Red dotted lines indicate infarct core. Scale bar 5 mm. B) Representative images of layers 2/3 of the motor cortex in WT male mice at 2 hps showing GluA2 AMPAR subunit (magenta) and VGluT1 (green) immunostaining. Scale bar 5 µm. C) Quantification of synaptic GluA2 puncta normalized to WT sham, total puncta of GluA2 normalized to WT sham, and total puncta of VGluT1 normalized to WT sham (n = 4). D,E) Same as B and C in the CA1 of hippocampus (n = 4). F) Overview of coronal sections of WT mouse at 1 day post-surgery (dps) in the motor cortex (top) and hippocampus (bottom), DAPI. The white dashed line delimits the infarct core. Scale bar 1mm. G) Representative images of the motor cortex (mCTX, top) and hippocampus (HC, bottom) in WT male mice at 1 dps showing DAPI (blue) and NeuN (green) immunostaining with *in situ* staining of TUNEL (yellow). Scale bar 500 µm. H) Quantification of TUNEL+ signal co-localizing with DAPI over total DAPI+ area in the ROI (%) (n = 4). I) Quantification of TUNEL+ signal colocalizing with NeuN over total NeuN+ area in the ROI (%) (n = 4). Bar graphs represent mean ± SEM, with individual data points per n. All statistics by unpaired two-tailed t-test, p-values on the graph.

At 2 hps, the motor cortex peri-infarct region (within 150 µm from the border of the injury) showed no significant change in total GluA2 levels, or co-localization of GluA2 puncta with the presynaptic marker VGluT1, compared to sham-operated mice immediately after ischemic injury (Figure 1B – C). In contrast, the hippocampal peri-infarct region exhibited a significant reduction in both total GluA2 puncta and GluA2- VGluT1 co-localized puncta relative to sham, suggesting loss of GluA2-containing AMPARs (Figure 1D – E). Previous research has shown that reduction in GluA2 leads to increased apoptotic cell death, particularly at later time points when apoptotic mechanisms are exacerbated, known as delayed apoptosis^6,18^. We found that, in response to focal ischemic stroke, delayed apoptosis (at 1 day post-stroke, dps) was significantly higher in the hippocampus than in the motor cortex following focal ischemic stroke (Figure 1F – I, S1B). These findings demonstrate that early GluA2 alterations differ between selectively vulnerable brain regions, suggesting that regional factors acting upstream of neuronal GluA2 trafficking may contribute to differential susceptibility after ischemic stroke.

### Early astrocytic chordin-like 1 expression differs between brain regions after ischemic stroke

Our previous research demonstrated that astrocytes regulate GluA2 levels during synaptic maturation and in response to ischemic stroke in the cortex via Chrdl1^12,14^. We previously observed rapid changes in GluA2 levels in hippocampal neurons within the first 20 minutes of exposure to ischemic conditions, but not in cortical neurons^6^. This differential effect was recapitulated in focal ischemic stroke models targeting the motor cortex or the hippocampus (Figure 1). We previously demonstrated that endogenous Chrdl1 expression is highly heterogeneous across brain regions, with substantially higher expression in the upper layers of the cortex than in the hippocampus (Figure S2A)^12,14^. If this early stroke-induced GluA2 regulation is influenced by endogenous Chrdl1 expression, we reasoned that regional differences in GluA2 stability should be accompanied by corresponding changes in early Chrdl1 expression after stroke. To address this, we quantified Chrdl1 mRNA in peri-infarct astrocytes using single-molecule fluorescent *in situ* hybridization (smFISH) against Chrdl1 and the astrocyte marker Slc1a3^12,15^. At this early time point (2 hps), peri-infarct astrocytes in the motor cortex showed a significant increase in Chrdl1 expression (Figure 2A – C, S2B), whereas peri-infarct astrocytes in the hippocampus showed no statistically significant change compared to sham-operated controls (Figure 2D – F). In contrast, we previously showed that by 7 dps, Chrdl1 is significantly upregulated in the peri-infarct astrocytes in both brain regions^12,15^.

**Figure 2.**
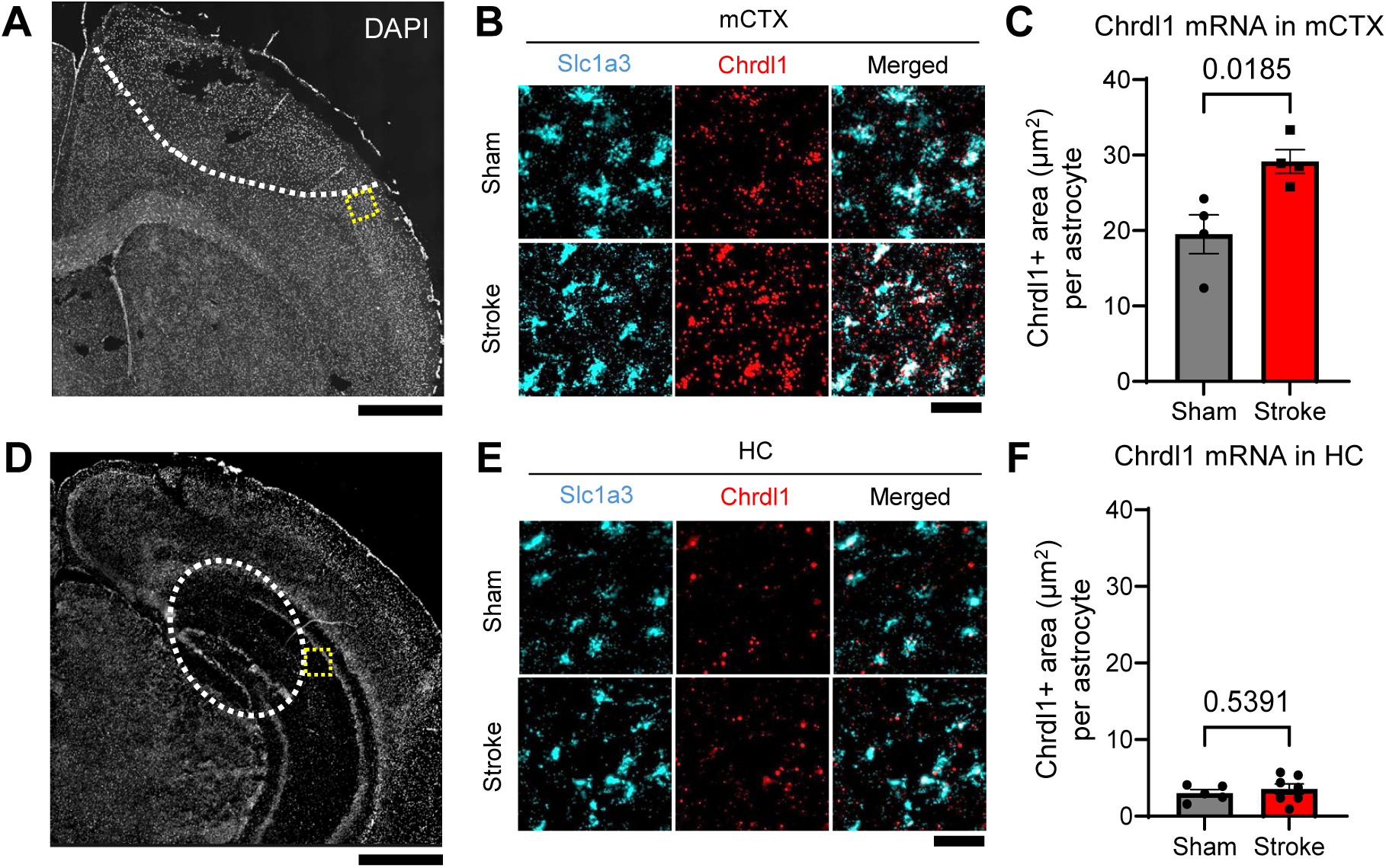
Early astrocytic chordin-like 1 expression differs between brain regions after ischemic stroke. A) Overview of coronal section of a WT mouse 2 hours after motor cortex (mCTX) stroke, visualized by DAPI. The white dashed line delimits the injury core, and the yellow box highlights the region of interest (ROI) imaged and analyzed. Scale bar 1 mm. B) Representative images of Slc1a3 (cyan) and Chrdl1 (red) single-molecule fluorescent *in situ* hybridization (smFISH) in the upper layers of the motor cortex in the peri-infarct area of male WT mice at 2 hps. Scale bar 50 µm. C) Quantification of Chrdl1+ area per astrocyte in the upper layers of the motor cortex in the peri-infarct region of male WT mice at 2 hps in sham and stroke mice (n = 4). D-F) Same as A, B, and C in the hippocampus in sham (n = 5) and stroke (n = 7) mice. Bar graphs represent mean ± SEM, with individual data points per mouse. All statistics by unpaired two-tailed t-test, p-values on the graph.

Taken together, these results suggest that Chrdl1 displays a distinct spatiotemporal expression profile following ischemic stroke, which may be leveraged to investigate region- and time-specific mechanisms underlying neuronal vulnerability and neuroprotection after stroke.

### Chordin-like 1 limits early GluA2 loss in hippocampal neurons under ischemic conditions

Having established that hippocampal neurons selectively lose GluA2 after ischemic injury (Figure 1D, E)^6^, and coinciding with the fact that peri-infarct astrocytes in the hippocampus do not upregulate Chrdl1 (Figure 2D – F), we next asked whether early induction of astrocytic Chrdl1 expression could preserve GluA2 levels under ischemic conditions, similar to the endogenous response observed in the more resilient motor cortex^12^. To test this, primary hippocampal neurons were treated with Chrdl1 prior to OGD. Surface and intracellular GluA2 levels were then quantified and compared to control cultures that were either not exposed to OGD or treated with vehicle (Figure 3A, S3A). As expected, vehicle-treated hippocampal neurons subjected to OGD significantly reduced surface GluA2 levels compared to non-OGD control cultures (Figure 3B – C). No differences in intracellular levels of GluA2 were detected under any condition. In contrast, Chrdl1-treated hippocampal neurons maintained stable surface GluA2 levels following OGD, comparable to non-OGD cultures (both vehicle- or Chrdl1-treated, Figure 3B – C). This phenomenon was observed immediately after OGD (Figure 3B – C) but not at later time points following return to glucose- and oxygen-containing media for 3 days (referred to as reperfusion, Figure 3D – E).

**Figure 3.**
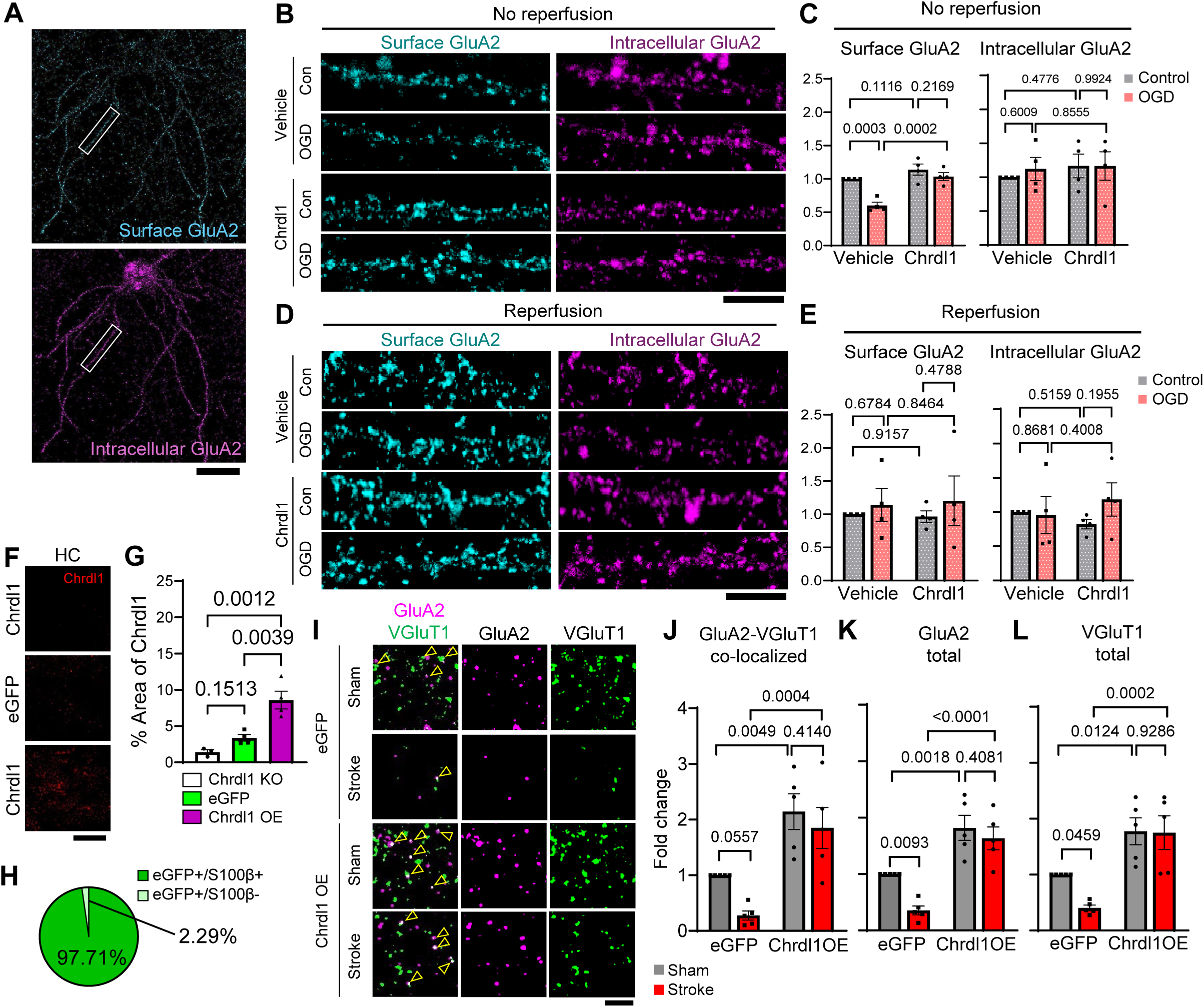
Chordin-like 1 limits early GluA2 loss in hippocampal neurons under ischemic conditions. A) Confocal images of whole cultured hippocampal neurons stained against GluA2 using a two-step antibody feeding protocol for surface (cyan) or intracellular (magenta) GluA2. White box indicating secondary dendrite length used for analysis. Scale bar 25 µm. B) Representative images of neurons treated with Chrdl1 or vehicle after 20 min of OGD and no reperfusion. Scale bar 5 µm. C) Surface and intracellular GluA2 were quantified on secondary hippocampal dendrites (ROI length: 25 µm). For each condition, we analyzed 3 – 4 dendrites per neuron, selecting at least three neurons across three coverslips. Data represents four independent cell culture preparations (n = 4). D – E) Same as B – C but neuronal cultures were incubated in normoxia and glucose-containing media for 72 h (reperfusion). Bar graphs represent mean ± SEM, with individual data points representing different cell cultures. Statistics by two-way ANOVA, followed by uncorrected Fisher’s LSD multiple-comparisons test and p-values on the graph. F) Representative images of AAV-eGFP (eGFP, AAV-PHP.eB::GfaABC1D-eGFP-4x6T) and AAV-Chrdl1OE (Chrdl1 OE, AAV-PHP.eB::GfaABC1D-Chrdl1-HA-4x6T) mice at 2 weeks after viral injection. Immunohistochemistry against Chrdl1 (red) in the CA1 of the hippocampus. Scale bar 25 µm. G) Quantification of Chrdl1 as % area positive for Chrdl1 signal in the CA1 hippocampus in Chrdl1 KO (n = 3), eGFP (n = 4), and Chrdl1 OE (n = 4) mice. Bar graphs represent mean ± SEM, with individual data points per mouse. Statistics by one-way ANOVA, followed by Holm-Šídák test, p-values on the graph. H) Quantification of % eGFP+ cells co-expressing or not S100β- in the CA1 of the hippocampus 2 weeks after AAV injection (n = 3). I) Representative images of CA1 region of hippocampus from mice that were injected with AAV-eGFP (eGFP, control) or AAV-Chrdl1 to overexpress Chrdl1 in astrocytes (Chrdl1 OE) at 2 hps showing GluA2 AMPAR subunit (magenta) and VGluT1 (green) immunostaining. Scale bar 5 µm. J) Quantification of GluA2 and VGluT1 colocalized puncta, K) total GluA2 puncta and L) total VGluT1 puncta (n = 5). Bar graphs represent mean ± SEM, with individual data points per mouse. Statistics by two-way ANOVA, followed by uncorrected Fisher’s LSD multiple-comparisons test and p-values on the graph.

Based on these observations, we asked whether astrocytic overexpression of Chrdl1 *in vivo* prior to the ischemic stroke could preserve GluA2 levels in the hippocampus following focal ischemic stroke in that brain region. Chrdl1 expression in the hippocampus under physiological conditions is low compared to cortical upper layers^12,14^ (Figure S2A), and early stroke-induced upregulation of Chrdl1 is absent in the hippocampus (Figure 2D – F), occurring only at later time points (7 dps)^12^. To this end, we generated an AAV to overexpress Chrdl1 specifically in astrocytes (AAV-PHP.eB::GfaABC1D-Chrdl1-HA-4x6T, a.k.a. Chrdl1 OE), along with a control AAV expressing eGFP (AAV-PHP.eB::GfaABC1D-eGFP-4x6T, a.k.a. eGFP) (Figure S3B – H). Both constructs were packaged in the PHP.eB capsid to enable systemic administration^19,20^ and avoid intracerebral injections that could induce secondary traumatic injury and confound interpretation. As expected, mice administered Chrdl1 OE showed a significant increase in Chrdl1 expression, whereas mice receiving control eGFP showed no change (Figure 3F – H). Chrdl1 OE resulted in approximately 2-fold increase in GluA2-VGluT1 co-localized puncta (Figure 3I, J), total GluA2 puncta (Figure 3I, K) and VGluT1 puncta (Figure 3I, L) compared to eGFP controls, with no loss in response to stroke (Figure 3I – L). This was accompanied by a significant increase in GluA2 puncta not co-localized with VGluT1, used here as a proxy for extrasynaptic GluA2 localization (Figure S3J).

Together, these findings demonstrate that astrocytic Chrdl1 expression prior to the onset of the ischemic conditions promotes GluA2 recruitment, and stabilization after ischemic injury, limiting early stroke-induced loss of GluA2 typically observed in vulnerable hippocampal neurons.

### Early astrocytic Chordin-like 1 expression reduces delayed apoptotic neuronal death after ischemic conditions

Early loss of GluA2 is associated with increased neuronal vulnerability to ischemic injury and delayed apoptotic cell death and is considered a hallmark of acute brain injury and several neurodegenerative disorders^6,8^. We therefore tested whether Chrdl1-mediated preservation of GluA2 reduced apoptotic cell death under ischemic conditions. We first assessed apoptotic neuronal death in primary hippocampal neurons treated with Chrdl1 or vehicle, immediately after OGD or after reperfusion. Apoptosis, measured by immunocytochemistry against cleaved caspase 3 (CC3), a canonical marker of apoptosis, was unchanged immediately after OGD in all conditions (Figure 4A, C). However, the proportion of CC3+ cells was significantly increased in vehicle-treated cultures after reperfusion, an effect that was not observed in the Chrdl1-treated cultures (Figure 4B, D), indicating that Chrdl1 treatment prevented delayed neuronal death *in vitro*.

**Figure 4.**
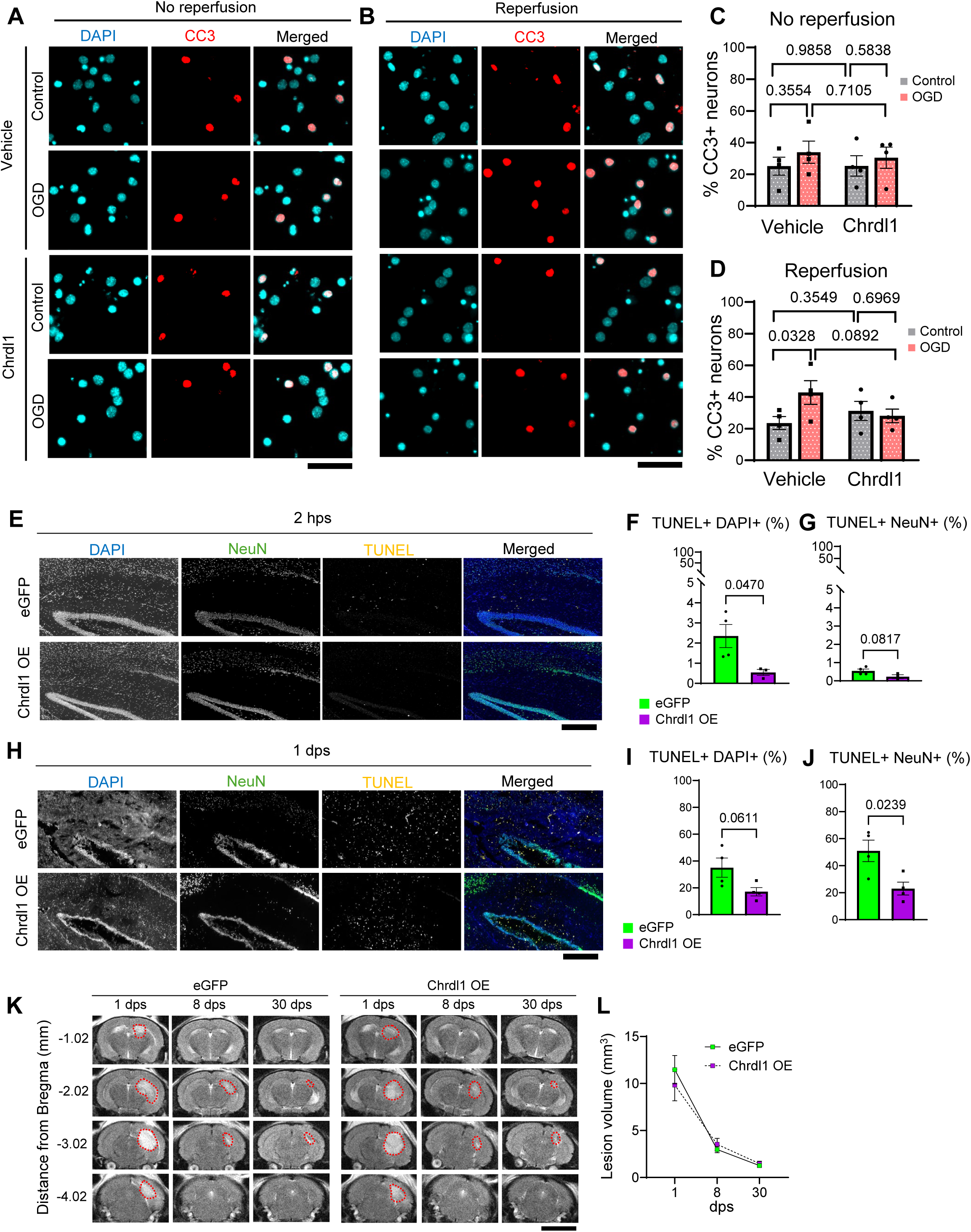
Early astrocytic Chordin-like 1 expression reduces delayed apoptotic neuronal death after ischemic conditions. A) Representative images of apoptotic cell death marker, cleaved caspase-3 (CC3) (red), and DAPI (cyan) of Chrdl1- or vehicle-treated hippocampal neurons after 20 min of OGD. Scale bar 25 µm. B) Same as A but neuronal cultures were returned to normoxia and glucose-containing media for 72 h after OGD (reperfusion). Scale bar 25 µm. C) Quantification of cell death, calculated as the percentage of CC3-positive nuclei relative to total DAPI-stained nuclei. Data represents four independent cell culture preparations (n = 4), with three coverslips analyzed per preparation and at least three regions of interest (ROIs) imaged per coverslip. D) Same as C but after reperfusion. Bar graphs represent mean ± SEM, with individual data points representing individual cell culture preparations. Statistics by two-way ANOVA, followed by uncorrected Fisher’s LSD multiple-comparisons test and p-values on the graph. E) Representative images of TUNEL (yellow), NeuN (green), and DAPI (blue) staining from the injury core of eGFP or Chrdl1 OE mice at 2 hps. Scale bar 500 µm. F) Quantification of TUNEL signal colocalizing with DAPI over total DAPI+ area in the ROI (%) of eGFP (n = 4) or Chrdl1 OE (n = 3) mice. G) Quantification of TUNEL- signal colocalizing with NeuN over total NeuN+ area in the ROI (%) of eGFP (n = 4) or Chrdl1 OE (n = 3) mice. H-J) Same as E – G but at 1 dps of eGFP (n = 4) and Chrdl1 OE (n = 3) mice. Bar graphs represent mean ± SEM, with individual data points per mouse. Statistics by unpaired two-tailed t-test p- values on the graph. K) Representative images of coronal T2-weighted MRI scans spanning from -1.02 mm caudal to -4.02 mm rostral to bregma of a male eGFP or Chrdl1 OE mouse at 1, 8, and 30 dps. Red dotted lines demarcate lesion. Scale bar 5 mm. L) Quantification of the lesion volume (mm^3^) in male eGFP (n = 11) or Chrdl1 OE (n = 11) mice. Statistics by two-way ANOVA, followed by Holm-Šídák test.

To determine whether this neuroprotective effect also occurred *in vivo,* we performed TUNEL labeling in brain sections from mice expressing either eGFP (control) or Chrdl1 OE following focal hippocampal stroke. TUNEL labeling was combined with DAPI and immunofluorescence against NeuN (neuronal marker). Co- localization of TUNEL and DAPI was used to quantify the proportion of apoptotic cells, whereas co- localization of TUNEL and NeuN was used to determine the proportion of apoptotic neurons in the hippocampus at 2 hps and 1 dps. As expected, apoptosis was minimal at 2 hps (Figure 4E – G). However, by 1 dps, mice overexpressing Chrdl1 in astrocytes exhibited lower apoptotic cell death (Figure 4H, I) with a significantly lower proportion of apoptotic neurons (Figure 4H, J), quantified as co-localized TUNEL and NeuN signal, compared to eGFP controls. Despite the reduction in delayed apoptotic neuronal death, MRI revealed no differences in lesion volume between Chrdl1 OE and eGFP mice at 1, 8, or 30 dps (Figure 4K, L).

Together, these results indicate that astrocytic overexpression of Chrdl1 prior to the onset of ischemic conditions reduces delayed apoptotic neuronal death without altering infarct volume.

### Early astrocytic Chordin-like 1 expression preserves hippocampal function but does not improve chronic recovery

Our results so far indicate that early Chrdl1 expression in astrocytes preserves GluA2 and limits delayed apoptotic neuronal death (Figure 3, 4). We next asked whether these effects translated into improved hippocampal function following ischemic stroke. To this end, we performed a longitudinal behavioral assessment spanning the acute (1 dps), subacute (3 – 8 dps), and chronic (≥30 dps) phases of recovery, evaluating hippocampal-dependent functions, including learning and memory, anxiety-like behavior, locomotion, and exploratory activity^21^ (Figure S4A).

First, using the open field test to assess anxiety-like behavior and exploration^22^, we found that Chrdl1 OE mice spent significantly less time in the center of the arena than the eGFP control mice during the acute phase, maintaining a behavioral profile more similar to baseline (Figure 5A). Although mice showed no differences in average speed throughout the study (Figure S4B), we detected other behavioral changes. During the acute phase, eGFP control mice did not show significant changes in distance traveled and time immobile, although Chrdl1 OE mice showed increased time immobile compared to the control mice at 1 dps (Figure 5A). By the subacute phase, Chrdl1 OE mice spent slightly less time in the center compared to eGFP mice, but differences in distance traveled were not observed between groups. However, eGFP mice displayed a significant increase in time immobile that was absent in Chrdl1 OE mice (Figure 5B). That difference was no longer evident during the chronic phase (Figure 5C), indicating that the behavioral effects of early Chrdl1 overexpression in astrocytes were transient. Overall, overexpression of astrocytic Chrdl1 seemed to preserve the baseline phenotype.

**Figure 5.**
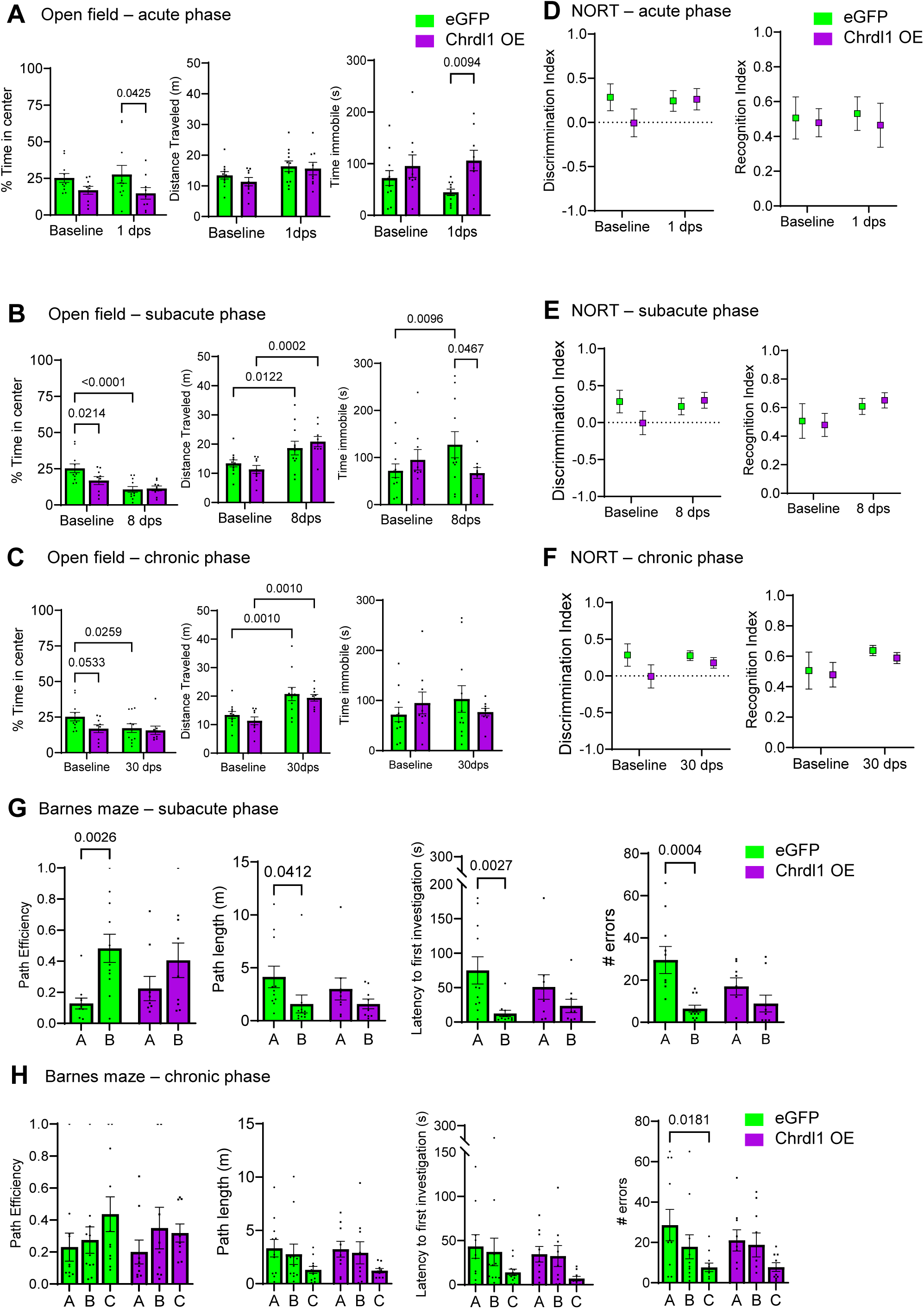
Early astrocytic Chordin-like 1 expression preserves hippocampal function but does not improve chronic recovery. A) Open field behavior testing in the acute phase (1 dps), B) subacute phase (8 dps), and C) chronic phase (30 dps) of eGFP (n = 11) and Chrdl1 OE (n = 9) mice compared to baseline testing with quantification of % time in center, total distance traveled (m), and time immobile (s). D) Discrimination and recognition indexes for the novel object recognition test (NORT) in the acute phase (1 dps), E) subacute phase (8 dps), and F) chronic phase (30 dps) of eGFP (n = 11) and Chrdl1 OE (n = 8 – 9) mice compared to baseline . G) Barnes maze behavior testing on probe day in the subacute phase (7 dps) on eGFP (n = 11) and Chrdl1 OE (n = 9) mice with analysis of path efficiency, path length (m), latency (s), and number of errors to first locate either the previously trained baseline location A or subacute target location B. H) Same analysis during the chronic phase (36 dps) on eGFP (n = 11) and Chrdl1 OE (n = 9) mice analyzed with target locations A, B and C. Bar graphs represent mean ± SEM, with individual data points per mouse. Statistics by two-way ANOVA followed by uncorrected Fisher’s LSD multiple- comparisons test (A – G) or Holm-Šídák test (H) with only significant p-values (p < 0.05) on the graphs.

To assess object recognition memory, we used novel object recognition test (NORT)^23^. Chrdl1 OE mice and eGFP controls did not show significant differences at baseline or during the acute phase at 1 dps (Figure 5D). Differences during the subacute phase or the chronic phase were not detected in either group (Figure 5E, F), suggesting that the unilateral ischemic lesion was insufficient to promote deficits. Latency to explore the NO was similar between groups throughout the experiment (Figure S4C).

To further evaluate hippocampal-dependent cognition, we assessed spatial memory, spatial re-learning, retrieval, and cognitive flexibility using the Barnes maze at multiple time points with relocated escape positions. Testing was conducted before stroke (baseline) and during the subacute (4 – 8 dps) and chronic phases (31 – 36 dps). Because of the experimental design, mice could not be tested during the acute phase. During the subacute probe trial, Chrdl1 OE mice retained memory of the pre-stroke escape location (A) while simultaneously remembering the new escape location (B), as indicated by comparable path efficiency toward both locations, as well as latency to investigate either location and number of errors (Figure 5G). In contrast, eGFP mice showed significantly reduced path efficiency toward the pre-stroke location (A), with increased path length, latency and number of errors to location A (Figure 5G). This advantage observed in Chrdl1 OE mice was no longer present during the chronic phase (Figure 5H). Although some significant differences were observed between groups during training trials, the trends were similar, including path efficiency, path length, latency to reach the escape box, and number of errors, at any post-stroke time point (Figure S4D – F), with the exception of a small but significant difference in the number of errors at day 3 (Figure S4F).

Overall, these findings indicate that astrocytic Chrdl1 overexpression transiently preserves hippocampal-dependent function during the early stages of recovery after ischemic stroke, particularly spatial memory, whereas these benefits are not maintained during the chronic phase.

#### Loss of Chrdl1 induces shared synaptic proteomic changes across brain regions

Because endogenous Chrdl1 expression is substantially higher in the cortex than in the hippocampus^12,14^ (Figure 2, S2A), we next asked whether endogenous Chrdl1 contributes differently to the molecular organization of excitatory synapses in these two brain regions under physiological conditions. Establishing these baseline differences could provide insight into the regional context in which this neuroprotective mechanism is engaged following ischemic injury. To address this question, we isolated synaptosomes and cytosolic fractions from the cortex and hippocampus of adult male WT and Chrdl1 global KO mice^14^ under basal conditions and performed LC-MS/MS (Figure S5A, B). We first compared the overall proteomic impact of Chrdl1 deletion across brain regions and subcellular fractions, and found that, as expected, Chrdl1 loss had a much larger impact on the cortical proteome than in the hippocampal proteome, particularly in the cytosolic fraction (Figure 6A). Principal component analysis (PCA) of all samples revealed clear segregation according to brain region, subcellular fraction, and genotype, supporting the robustness of the dataset (Figure S5C). Given our focus on the synaptic functions of Chrdl1, we next concentrated our analyses on the synaptosome fraction.

**Figure 6.**
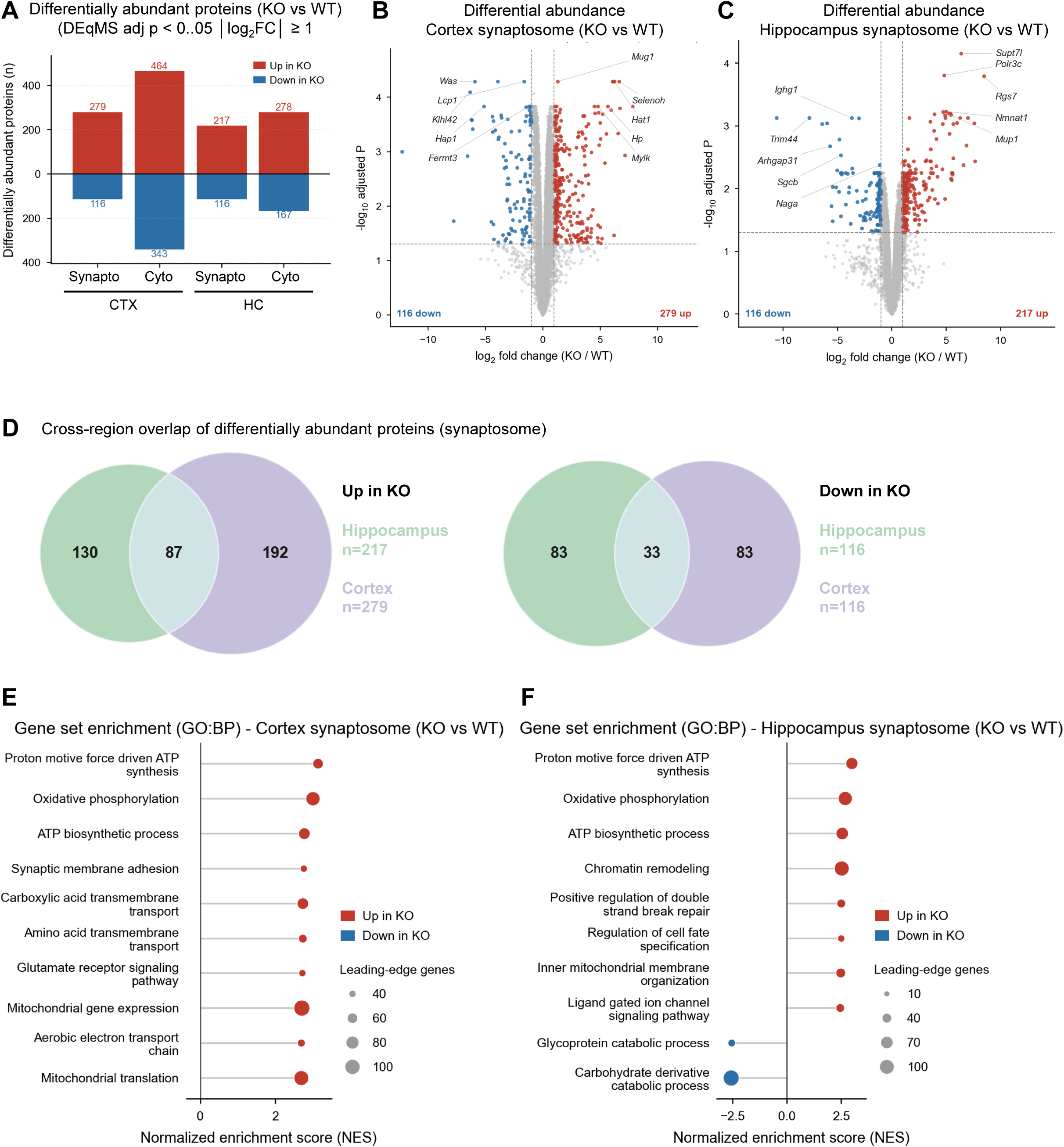
Loss of Chrdl1 induces shared synaptic proteomic changes across brain regions. A) Differentially abundant proteins in Chrdl1 KO vs. WT samples of cortical (CTX) and hippocampal (HC) synaptosomes and cytosol from adult mouse brain tissue (n = 3 per condition). B – C) Volcano plot of differentially abundant proteins comparing WT and Chrdl1 KO CTX synaptosomes or HC synaptosomes, respectively (n = 3 biological replicates per condition; fold changes are expressed as KO relative to WT). D) Analysis of uniquely and shared upregulated and downregulated proteins in CTX and HC synaptosomes shown via Venn diagrams. E) Gene set enrichment analysis (GSEA) of GO biological process terms in the CTX synaptosome and F) HC synaptosome comparisons. Proteins were ranked by the DEqMS moderated t-statistic, and enrichment was assessed for GO biological process gene sets from the mouse MSigDB collection (v2026.1.Mm)

Differential abundance analysis identified more altered proteins in cortical than in hippocampal synaptosomes, consistent with the overall proteomic comparison (Figure 6B, C, S5D, E). Comparison of differentially abundant proteins between brain regions revealed a substantial overlap, with 87 proteins increased and 33 proteins decreased in both cortical and hippocampal synaptosomes following Chrdl1 deletion, alongside a large number of region-specific changes (Figure 6D). These findings indicate that loss of Chrdl1 engages both shared and region-specific synaptic proteomic programs.

To determine whether these overlapping and region-specific protein changes converged on common biological processes, we performed gene set enrichment analysis (GSEA). Despite the larger proteomic remodeling observed in the cortex, both cortical and hippocampal synaptosomes showed enrichment of pathways related to glutamatergic signaling and mitochondrial function, including oxidative phosphorylation, ATP biosynthesis, and proton motive force-driven ATP synthesis (Figure 6E, F). These shared enrichments indicate that loss of Chrdl1 engages largely conserved synaptic pathways across brain regions despite marked differences in the magnitude of the proteomic response. Examination of the leading-edge proteins driving these enrichments revealed that glutamatergic signaling in both regions was supported by multiple AMPAR- and postsynaptic-associated proteins, including Gria1, Gria2, Gria3, Shank1, and Shank3 in the cortex, and additional kainate and NMDA receptor subunits in the hippocampus (Figure S5F). Although Gria2 was among the leading-edge proteins contributing to glutamatergic signaling in both regions, it was not identified as differentially abundant in the differential abundance analysis. Similarly to the glutamatergic signature, mitochondrial pathways were driven by coordinated increases in proteins involved in oxidative phosphorylation and electron transport chain function in both brain regions.

Together, these findings demonstrate that Chrdl1 deletion produces more extensive synaptic proteomic remodeling in the cortex than in the hippocampus while converging on common glutamatergic and mitochondrial pathways in both regions.

## DISCUSSION

The key finding of our study is that a developmental astrocyte-derived mechanism can be repurposed following ischemic stroke to regulate neuronal vulnerability. We show that the astrocytic protein Chrdl1, previously characterized as a regulator of synaptic maturation during development^14^, preserves GluA2- containing AMPARs and limits delayed apoptotic neuronal death when present during the hyperacute phase (<1 dps) of ischemic injury. Interestingly, this protective mechanism appears to depend on both the brain region and the timing of Chrdl1 expression in astrocytes. While endogenous Chrdl1 is rapidly induced in cortical peri-infarct astrocytes after a focal stroke in that brain region, this early response is absent in the hippocampus, a more vulnerable brain region^6,24,25^. Increasing early Chrdl1 expression in astrocytes was sufficient to preserve GluA2, reduce delayed apoptotic neuronal death in the hippocampus, and transiently improve hippocampal function. Together, these findings suggest that developmental astrocyte-derived mechanisms can be repurposed following ischemic injury, but that their protective effects depend on when and where they are engaged.

The temporal profile of Chrdl1 expression in astrocytes appears to be critical to its function after focal ischemic stroke. During the acute post-stroke stage, excessive glutamate release and excitotoxic signaling drive rapid loss of GluA2-containing AMPARs and promotes apoptotic neuronal death^26–29^. Under these conditions, early Chrdl1 expression preserves GluA2, as observed in the motor cortex^12^. However, this protective effect was not maintained during the later post-stroke stages of recovery, when functional restoration relies on synaptic remodeling and plasticity rather than acute neuroprotection. Our findings suggest that sustained astrocytic Chrdl1 expression throughout recovery, may limit plasticity mechanisms that are normally engaged during these later post-stroke stages to restore neuronal connectivity and function^4,30,31^, consistent with its role during synaptic maturation in development.

Our findings support a model in which Chrdl1 promotes neuronal resilience by stabilizing GluA2-containing AMPARs during the acute phase of ischemic injury. Although previous studies have demonstrated that ischemic-like conditions promote rapid internalization of GluA2-containing AMPARs in hippocampal neurons and not in cortical neurons^6,7,13^, the mechanisms regulating this regional difference have remained unclear. Here, we show that increased early Chrdl1 expression may be responsible for preserving GluA2 levels both *in vitro* and *in vivo*, supporting the idea that astrocyte-derived signals contribute to differential regulation. The predominant AMPAR heteromer found in hippocampus is GluA2/GluA1^32^, and previous studies have shown that ischemic preferentially promotes endocytosis and degradation of GluA2/GluA3- AMPARs^13^. One possibility is that Chrdl1 shifts GluA2-containing AMPARs away from degradation and toward recycling and membrane stabilization. Although this mechanism remains to be directly tested, it is consistent with our observation that Chrdl1 increases GluA2 levels without any differences in intracellular GluA2.

Our behavioral findings further support the idea that the effects of Chrdl1 depend on the stage of stroke recovery. Increasing Chrdl1 expression prior to the insult resulted in a transient functional benefit after hippocampal ischemic stroke, most evident as preservation of spatial memory during the subacute phase. This benefit was not sustained during chronic recovery, consistent with a time-dependent role for Chrdl1 in which early neuroprotection does not translate into improved long-term functional recovery. This temporal profile is consistent with the established role of Chrdl1 during development, where it limits experience- dependent plasticity^14^. We previously showed that deletion of Chrdl1 improved long-term functional recovery following motor cortex focal stroke^12^, suggesting that sustained Chrdl1 expression may become maladaptive and restrict synaptic remodeling. Interestingly, in that study, Chrdl1 KO mice, which expressed ∼20% of endogenous Chrdl1 expression^14^, also showed reduced apoptotic cell death within the cortical injury core at 1 dps^12^, whereas here we show that increasing Chrdl1 in the hippocampus reduced delayed apoptotic neuronal death. These apparently opposing findings may be a consequence of the significantly different endogenous expression of Chrdl1 in the two brain regions. Chrdl1 is highly expressed in cortical astrocytes and rapidly upregulated after ischemic stroke, whereas its endogenously low and its early post-stroke upregulation is absent in the hippocampus. Thus, deleting Chrdl1 from a high-expressing region versus increasing its expression in a low-expressing region may not represent equivalent opposing manipulations, but instead suggest that neuronal protection depends on appropriate Chrdl1 signaling within the regional context. Together with our current findings, these observations reinforce our model in which Chrdl1 expression may promote neuroprotection early after ischemic stroke by stabilizing GluA2-containing AMPARs but becomes maladaptive in the long-term by limiting plasticity mechanisms required for long-term recovery. Our data suggest that the endogenous spatiotemporal expression of Chrdl1 in astrocytes determines whether, and to what extent, this mechanism is engaged following ischemic injury.

The proteomic analysis further supports the idea that Chrdl1 regulates a conserved synaptic program across brain regions. Although endogenous Chrdl1 deletion altered the cortical synaptic proteome more extensively than the hippocampal proteome, glutamatergic signaling and mitochondrial function emerged as shared biological themes in both regions. Notably, despite the enrichment of glutamatergic signaling pathways, neither our previous RNA-sequencing analysis^14^ nor the present synaptosomal proteomics identified a reduction in overall Gria2 expression in the Chrdl1 KO, supporting the idea that Chrdl1 primarily regulates the synaptic localization or stabilization of GluA2-containing AMPARs rather than their transcription or overall abundance. These observations are consistent with the markedly higher physiological expression of Chrdl1 in cortical astrocytes and suggest that regional differences arise from the degree to which synapses depend on endogenous Chrdl1 rather than from fundamentally distinct molecular mechanisms. Importantly, our observation that increasing Chrdl1 expression in the hippocampus promotes GluA2 and limits neuronal death indicates that this developmental program remains available in hippocampal neurons despite the low endogenous expression of Chrdl1.

Our study positions astrocytes as dynamic regulators of the balance between neuroprotection and adaptive plasticity, highlighting that optimal recovery after ischemia may depend not only on preventing early excitotoxic injury but also on permitting controlled synaptic remodeling during later phases. Our studies suggest that the spatiotemporal engagement of developmental mechanisms, such as Chrdl1 signaling, is a key determinant of neuronal vulnerability and functional recovery after ischemic stroke.

### Limitations of the study

One limitation of this study is that we did not directly determine whether Chrdl1-induced increase and stabilization of GluA2-containing AMPARs altered Ca^2+^ permeability or synaptic function. Previous studies have demonstrated that ischemia and OGD promote the removal of GluA2-containing AMPAR from hippocampal neurons, resulting in the synaptic expression of GluA2-lacking, Ca^2+^-permeable AMPARs that contribute to toxic Ca^2+^ influx and delayed apoptotic neuronal death^6,7,9,13,33^. Consistent with this model, Chrdl1 preserved surface GluA2 *in vitro* and reduced CC3 expression after OGD, while astrocytic Chrdl1 overexpression reduced TUNEL labeling *in vivo* after hippocampal focal stroke. These findings support the interpretation that Chrdl1 may limit delayed apoptotic neuronal death by maintaining a higher proportion of GluA2-containing, Ca^2+^-impermeable AMPARs at the neuronal surface. Future electrophysiological or Ca^2+^ imaging studies will be needed to directly test this mechanism.

## RESOURCE AVAILABILITY

### Lead contact

Requests for further information and resources should be directed to and will be fulfilled by the lead contact, Dr. Elena Blanco-Suárez.

## Materials availability

Plasmids generated in this study have been deposited to Addgene, pAAV-GfaABC1D-eGFP-4x6t (#260900) and pAAV-GfaABC1D-Chrdl1-HA-4x6t (#260901). Additional materials that are not commercially available may be requested directly from the lead contact.

### Data and code availability

#### Data

All data reported in this paper will be shared by the lead contact upon request. The data used for analysis are available in the Mendeley Data repository (doi: 10.17632/6bgtkrv24f.1). The proteomics data are available in the PRIDE repository under accession number PXD082639.

#### Code

The code used in this study is publicly available on Zenodo at https://doi.org/10.5281/zenodo.21924869.

#### Additional information

Any additional information required to reanalyze the data reported in this paper is available from the lead contact upon request.

## Supporting information

Table S1 Statistics

## ACKNOWLEDGMENTS

This work has been funded by the American Heart Association (AHA) Career Developmenta Award, and Bridge Funding Awards to EB-S (award numbers 937695, 26BCDA1622714 and 26BIPA1622640) and the AHA Predoctoral Fellowship to BRB (award number 24PRE1198593). We are grateful to The Wistar Institute’s Proteomics & Metabolomics Shared Resource for providing technical support. Funding support for The Wistar Institute core facilities was provided by Cancer Center Support Grant P30 CA010815.

## AUTHOR CONTRIBUTIONS

Conceptualization EB-S; methodology, BRB, RBH, AJG, STC, and EB-S; Investigation, BRB, RBH, AP, AJG; writing—original draft, review & editing, BRB and EB-S; funding acquisition, EB-S; resources, EB-S and STC; supervision, EB-S.

## DECLARATION OF GENERATIVE AI AND AI-ASSISTED TECHNOLOGIES

During the preparation of this work, the authors used ChatGPT (GPT-5.5, OpenAI) solely to improve grammar and language. The authors reviewed and edited all AI-generated suggestions as needed and take full responsibility for the content of this publication.

## METHOD DETAILS

### Animals

All animal work was approved by the Institutional Animal Care and Use Committee (IACUC) of Thomas Jefferson University. All mice were housed in the Thomas Jefferson University Animal Facility at the Bluemle Life Sciences Building under a 12 h light–dark cycle with controlled temperature and humidity and had *ad libitum* access to food and water. All animal experiments were conducted in accordance with ARRIVE guidelines. Male wild-type (WT) (C57BL/6J, RRID: IMSR_JAX:000664) and Chrdl1 KO mice^14^ (the latter only for preparation of synaptosomes and cytosolic fractions) at 3 – 5 months of age were used. WT mice weight ranged between 25 – 30 g with no significant weight loss after surgeries or retro-orbital injections.

Embryonic hippocampal tissue used for primary neuronal cultures was gifted by Dr. Le Ma at Thomas Jefferson University. In summary, hippocampal tissue was obtained from rat embryos at embryonic day 17 – 18 (E17 – 18), from timed-pregnant Long–Evans rats (Charles River Laboratories; RRID: RGD_2308852). Dams were euthanized by CO₂ inhalation, in accordance with Dr. Ma’s IACUC approved protocol and following ARRIVE guidelines. Rat embryos were collected from Dr. Le Ma’s laboratory, maintained in Leibovitz’s L-15 medium (Thermo Fisher Scientific Cat# 11415064) and transferred to our laboratory.

### Primary hippocampal neuron al cultures and treatments

E18 rat embryos in Leibovitz’s L-15 medium (Thermo Fisher Scientific Cat# 11415064) were used for hippocampal dissection in HBSS 1X (Thermo Fisher Scientific Cat# 14170112) supplemented with 1 mM HEPES (Thermo Fisher Scientific Cat# 15630-080). Hippocampi from 7-8 embryos were washed gently 3 times with filtered HBSS (Thermo Fisher Scientific 14170112) and 1 mM HEPES (Thermo Fisher Scientific 15630-080) and incubated with 0.25% trypsin (Thermo Fisher Scientific 25200-056) for 15 min at 37 °C, with gentle swirls every 5 min. Tissue was gently washed three times with 10% FBS (Thermo Fisher Scientific A52567-01) and culture media. The culture medium was sterile-filtered neurobasal medium (Thermo Fisher Scientific 21103049) supplemented with 50x B27 (Thermo Fisher Scientific 17504044), 100x GlutaMAX (Thermo Fisher Scientific 35050061), and Penicillin-Streptomycin 100x (Thermo Fisher Scientific 15070063). Tissue was gently triturated with a fire-polish glass pipette. Neurons were then plated at low density of about 80,000 per coverslip (Fisher Scientific 12541000) on 12mm-diameter circular coverslips (Fisher Scientific 12541000) previously coated with 0.1 mg/ml PDL (Thermo Fisher Scientific A38904-01) and 15 µg/ml laminin (Thermo Fisher Scientific 23017-015) in a 24-well plate. mRNA from cultures were collected, as described later, to assess purity (Figure S2C). Cultures were treated at DIV 10 and 13 with either Chrdl1 (1 µg/ml in 0.1% BSA 4 mM HCl) (R&D 1808-NR/CF) or vehicle (0.1% BSA 4 mM HCl) (Sigma-Aldrich A4161) prior to OGD or control conditions.

### Photothrombosis in the motor cortex

Focal ischemic stroke was induced in the right motor cortex using the photothrombosis model, as previously described^12^. Adult male mice at 16 weeks of age were anesthetized with isoflurane (5% induction; 1.5–2% maintenance in 1 L/min O₂) and attached to a stereotaxic frame. Body temperature was monitored and maintained at 37 ± 0.5 °C using a feedback-controlled heating pad (Kent Scientific RT-JR). Fur over the scalp was removed, and the skin was disinfected with alternating swabs of 70% ethanol (Decon Labs 105118) and providone-iodine, 10% (Betadine 67618-150-17). Rose bengal (Fisher Scientific 632-69-9) at 10 mg/mL in sterile 0.9% NaCl prepared fresh, protected from light, and filtered through a 0.22 µm syringe filter, was administered via retro-orbital injection at a final dose of 50 mg/kg. A midline scalp incision was made to expose the skull, and bregma was identified as the stereotaxic reference point. The right hemisphere was targeted at +0.3 mm (AP), -1.5 mm (ML) from bregma. Five minutes after Rose Bengal injection, the skull over the target region was illuminated for 10 min using a diode laser (λ=520 nm, Thor Labs LP520-SF15) set to 10 mW with a 2 mm beam diameter, positioned 2 mm above the mouse skull. Following illumination, the skull was rinsed with sterile 0.9% NaCl, and the incision was closed using tissue adhesive (Vetbond 3M 1469Sb). Triple antibiotic ointment (Globe) and 5% lidocaine (Mohnark) were applied topically. Mice recovered on a heating pad for at least 30 min before being returned to their home cages. Sham-operated animals underwent identical procedures but received 0.9% NaCl retro-orbital injection instead of rose bengal.

### Photothrombosis in the hippocampus

Photothrombotic ischemia targeting the right hippocampus was performed using a fiber-coupled laser delivery approach following our published protocol^16^. Mice were anesthetized and surgically prepared as described for motor cortex photothrombosis. The laser diode was set at 7 mW, and attached to a FC/PC ferrule patch cable (Thor Labs M83L01) through the mating sleeve (Thor Labs ADAL1) connecting to a fiber optic cannula (Thor Labs CFMLC12L02). A burr hole was drilled (World Precision Instruments) using 0.45 mm drill bit (Fisher Scientific 10-001-085) at −2.7 mm (AP) and −2.2 mm (ML) from bregma. The optic fiber cannula (Thor Labs CFMLC12L02), attached to the stereotax, was lowered to −1.5 mm (DV) to position the fiber tip to reach the hippocampus. Rose bengal at a concentration of 5 mg/mL in sterile 0.9% NaCl; prepared fresh, filtered, and protected from light, was administered via retro-orbital injection at a final dose of 25 mg/kg. The target region was then illuminated for 10 min, and then the optic fiber was slowly withdrawn. The skull was rinsed with sterile saline (0.9% NaCl), and the incision was closed with Vetbond tissue adhesive. Triple antibiotic ointment and 5% lidocaine were applied topically. Sham-operated animals underwent identical procedures but received sterile saline (0.9% NaCl) retro orbital injection instead of rose bengal.

### Tissue collection and preparation

Brains were collected at 2 hours or 1 day after surgeries. Mice were anesthetized by intraperitoneal injection of Ketamine (100 mg/Kg, Mylan) and Xylazine (20 mg/Kg, Anased) and transcardially perfused with ice-cold PBS 1X only or PBS 1X followed by 4% PFA, depending on the following experimental procedures. Brains were collected, embedded in OCT (Scigen 4583) and stored at -80°C until further processing.

### Synaptic staining and puncta analysis on brain tissue

Fluorescence immunohistochemistry was performed on fresh frozen 16 µm brain coronal sections collected at 2 hps in a cryostat (Microm HM550) approximately sectioned between 1.18 mm and -0.62 mm or between

-3.28 mm and -2.8 mm AP from bregma to include the motor cortex or the hippocampus, respectively. Brain sections were collected on Superfrost Plus microscope slides (Fisher Scientific 22037246), and washed three times with PBS 1X to remove OCT. Sections were then fixed for 8 min in ice-cold methanol, washed 3 times with PBS 1X, and then fixed with cold 4% PFA for 8 min, followed by three PBS 1X washes. Next, hydrophobic barriers were drawn around tissue using a PAP pen (Fisher Scientific NC9204359). Sections were blocked with 5% normal goat serum (NGS) and 0.3% TritonX-100 in PBS 1X for 1 hour at room temperature (∼22°C – 24°C) in a humidified chamber. Sections were incubated in primary antibody anti- GluA2 (Sigma Millipore AB1768-I; RRID:AB_2922404) at 1:500 and anti-VGluT1 (Sigma Millipore AB5905; RRID:AB_2301751) at 1:1000 in antibody buffer (5% NGS, 0.3% Triton X-100, PBS 1X and 5 mM L-lysine) overnight at 4°C in a humidified chamber. Sections were washed the next day three times with PBS 1X and incubated in secondary antibody anti-rabbit Alexa Fluor 594 (Thermo Fisher Scientific A11037; RRID:AB_2534095) and anti-guinea pig Alexa Fluor 647 (Thermo Fisher Scientific A21450; RRID:AB_2535867) at 1:500 in antibody buffer for 2 h at room temperature (∼22°C – 24°C) in a humidified chamber, followed by three PBS 1X washes. Slides were mounted with Slowfade gold antifade mountant with DAPI (Thermo Fisher Scientific S36939) using coverslips 22 mm x 50 mm 1.5 thickness (Globe Scientific 141410) and sealed with clear nail polish.

The CA1 area of the hippocampus was imaged in the peri-infarct area (immediately adjacent to the border of the core of the injury) or the homologous region in the sham brains on a Leica SP5 confocal microscope (RRID:SCR_020233) using a 63x/1.4 NA oil-immersion objective as 8-bit images at 1024 X 1024 pixels (pixel size 120.1 x 120.1 nm Figure 1B, D, and pixel size 125 x 125 nm in Figure 3I, M), as a 3.02 µm z-stack of 10 steps (step size 0.34 µm, 10% overlap). Image analysis of synaptic puncta was conducted blind to treatment and as 3D images using IMARIS software (Oxford Instruments, Bitplane version 9.8.2, RRID:SCR_007370). Gaussian filter of 0.125 µm was applied to all channels to reduce noise. Using the spot detection function, puncta were defined in each channel as spheres of average diameter of 0.5 µm. Using the co-localization spots function, we were able to define GluA2 spots co-localized and non- colocalized with the VGluT1 spots. Co-localization was defined as proximity of the postsynaptic GluA2 spot to presynaptic VGluT1 spot of 0.5 µm or less from each other’s sphere center. All representative images are orthogonal projections of 3 contiguous steps from a z-stack.

### Generation of adeno-associated virus (AAV)

AAV::GfaABC1D-Chrdl1-4x6T plasmid was produced using Gibson assembly. The viral backbone was generated from AAV::GfaABC1D-smMyc-4x6T (Addgene plasmid # 196415; http://n2t.net/addgene:196415 ; RRID:Addgene_196415))^34^ using restriction enzyme digest (New England Biolabs). Isoform 1 of Chrdl1 was generated through a combination of overlapping plasmid cDNA for isoform 2 (TransOMIC Technologies BC066832) and a synthetic fragment specific to isoform 1 (Integrated DNA Technologies); these fragments were assembled with the viral backbone using HiFi Master Mix (New England Biolabs E2621L). Plasmids were sequenced using whole-plasmid sequencing (Plasmidsaurus). PHPeB::GfaABC1D-Chrdl1-4x6T virus was produced in accordance with the US NIH Guidelines for Research Involving Recombinant DNA Molecules and the University of California – Los Angeles Institutional Biosafety Committee. Virus was produced as previously described^20,34^. Briefly, HEK293T cells (ATCC CRL- 3216, RRID:CVCL_0063) were transiently transfected with transfer plasmid containing GfaABC1D-Chrdl1- 4x6T, capsid plasmid for PHP.eB, and helper plasmid. Viral particles were isolated from the media via polyethylene glycol precipitation and released from cells using salt-active nuclease (ArcticZymes 70910- 202). Virus was then purified using iodixanol column purification and quantified using qPCR directed against the WPRE. WPRE primers [5’→3’]: forward:, GGCTGTTGGGCACTGACAAT; reverse, CCGAAGGGACGTAGCAGAAG. pUCmini-iCAP-PHP.eB was a gift from Viviana Gradinaru (Addgene plasmid # 103005 ; http://n2t.net/addgene:103005; RRID:Addgene_103005).

The HA tag was inserted by polymerase chain reaction (PCR) using Phusion High-Fidelity DNA polymerase (Thermo F530S). Primers [5’→3’] forward: GAGACCTGAAAAGGACCACTGTTACCCTTACGATGGTACCGGATTACGCATAGGGATC; reverse: GAATGCCCATATGGGATCCCTATGCGTAATCCGGTACATCGTAAGGGTAACAGTGGT were designed using NEBuilder (New England BioLabs). To generate the control AAV expressing eGFP instead of Chrdl1-HA, the plasmid was cut by double restriction enzyme digest with PacI (NEB #R0547S) and BamHI-HF (NEB #R3136S) to remove Chrdl1. The eGFP sequence was PCR amplified using Q5 High-Fidelity DNA polymerase (NEB #M0515) using primers [5’→3’] forward: TCGTTCTTAATTAAATGGTGAGCAAGG; reverse, TTCTATCAGGATCCTTACTTGTACAGC designed using NEBuilder (New England BioLabs). The eGFP fragment was ligated to the viral backbone using T4 DNA Ligase (NEB M0202S). Both constructs were confirmed by whole-plasmid sequencing (Primordium Labs). Viral purification and titer was determined via ddPCR by Franklin Biolabs Vector Core.

### AAV delivery

AAVs were systemically delivered using retro-orbital injections. Mice were anesthetized with 5% inhaled isoflurane. The injections were performed using U-100, 3/10cc, 31 G X 5/16” needles. To perform retro-orbital injection, the needle was inserted into the medial canthus at a 45° angle behind the ocular globe. Virus was diluted in sterile saline to deliver a consistent viral dose per mouse. Injection volume was typically 50 µL and was increased up to 100 µL as needed to compensate for differences in viral titer. AAVs were delivered at 3 X 10^12^ vg/mouse. Petrolatum ophthalmic ointment (Systane) was applied to the injected eye afterwards to prevent discomfort.

### Immunostaining and analysis of eGFP or Chrdl1 on brain tissue

Fluorescence immunohistochemistry was performed on 16-µm coronal brain sections from mice perfused with PBS 1X followed by 4% PFA, and collected 2 weeks after AAV retro-orbital injection. Sections were washed three times in PBS 1X and blocked for 1 h at room temperature in blocking buffer in a humidified chamber. Primary antibodies were diluted in antibody buffer and incubated overnight at 4°C in a humidified chamber. Primary antibody combinations included anti-GFP at 1:1000 (Aves Labs, GFP-1020) with anti- S100β at 1:100 (Abcam, ab52642), and anti-Chrdl1 at 1:100 (MilliporeSigma, HPA000250) with anti-GFAP at 1:500 (Thermo Fisher Scientific, 13-0300). Sections were washed three times in PBS 1X and incubated for 2 h at room temperature with the corresponding Alexa fluor-conjugated secondary antibodies diluted in antibody buffer: goat anti-rabbit Alexa Fluor 594 at 1:500 (Invitrogen, A11037) with goat anti-chicken Alexa Fluor 488 at 1:200 (Invitrogen, A11039) or goat anti-rat Alexa Fluor 647 at 1:500 (Thermo Fisher Scientific, A21245). Sections were then washed three times in PBS 1X and mounted using SlowFade Gold antifade reagent with DAPI (Thermo Fisher Scientific S36939) using coverslips 22 mm x 50 mm 1.5 thickness (Globe Scientific 141410) and sealed with clear nail polish. Brain sections were imaged on a Leica Thunder Imager 3D Tissue using 20x/0.55 NA objective. Whole-brain tile scan images were taken as 16-bit tiles, at 31189 x 20329 pixels per tile (pixel size 0.610 x 0.610 µm) with a 10% overlap, and single 20x frames were taken as 16-bit images, at 2048 x 2048 pixels (pixel size 0.325 × 0.325 μm), 13.65 µm-thick z-stack of 10 steps with 10% overlap each step. A 700 × 600 µm ROI was selected in the ipsilateral hippocampus using consistent anatomical landmarks across all sections. eGFP+ cells and eGFP+/S100β+ cells were manually counted using the Cell Counter plugin in ImageJ. The percentage of eGFP+ cells co-expressing S100β was calculated. To determine Chrdl1 levels, a consistent threshold was applied to the 20x images to quantify % area positive for Chrdl1 signal.

### Single-molecule fluorescent *in situ* hybridization (smFISH) and analysis

Single-molecule fluorescent *in situ* hybridization (smFISH) was performed using RNAscope technology (ACD 323110) and following manufacturer instructions. PBS-perfused brains were cut in a cryostat into coronal brain sections of 16-µm thickness between 1.18 mm and -0.62 mm or between -3.28 mm and -2.8 mm relative to bregma for motor cortex and hippocampus, respectively, containing the injury core or the homologous brain region in sham brains. Brain sections were mounted on SuperFrost Plus microscope slides (Fisher Scientific 22037246) and stored at -20°C for next-day processing. Slides were gently rinsed 3 times in PBS 1X for 5 minutes, in gentle shaking, to remove any remaining OCT. Slides were fixed for 1 h at 4°C in cold 4% PFA made from EM grade 16% PFA ampules (EMS 15710) diluted in PBS 1X. Slides were washed twice with PBS 1X for 2 minutes in gentle shaking. Fixed slides were then dehydrated in sequential incubations in 50%, 70% and 100% ethanol for 5 min each. Brain sections were pre-treated with Protease IV (ACD 322336) for 16 minutes at room temperature. Sections were incubated with ACD probes targeting Chrdl1 (ACD, 442811-C1) and Slc1a3 (ACD, 430781-C2) for 2 h at 40°C. Following two washes in wash buffer 1X (2 min each, room temperature), probes were visualized using TSA Vivid fluorophores 570 (Channel 1) and 520 (Channel 2) (ACD, 323272 and 323271) according to the manufacturer’s instructions. We used the 3-plex negative probe (ACD 320871) for the negative control (Figure S2B). Slides were mounted with Slowfade gold antifade mountant with DAPI (Thermo Fisher Scientific S36939) using coverslips 22 mm x 50 mm 1.5 thickness and sealed with clear nail polish. Imaging was performed in the peri-infarct region or homologous region in the sham mice. Brain sections were imaged on a Leica Thunder Imager 3D Tissue using 20x/0.55 NA objective. Images were taken as 16-bit, 2048 x 2048 pixels (pixel size 0.325 × 0.325 μm), z-stack of 6.78 µm total thickness, made of five 1.69 µm steps with 10% overlap. Chrdl1 was imaged in channel 520 and Slc1a3 in channel 488. Using a semi-automatic custom-made macro for Image J^12,35^, Slc1a3 signal was used to determine ROIs, corresponding to individual astrocytes. A consistent threshold was applied to the Slc1a3 and Chrdl1 signal to quantify Chrdl1+ area (µm^2^) per astrocyte (Slc1a3+ cells).

### TUNEL and NeuN co-staining and analysis

4% PFA-perfused brains where coronally sectioned at 16-µm thickness. Fluorescence immunohistochemistry against NeuN (CST 24307; RRID:AB_2651140) at 1:500 dilution followed by incubation with secondary antibody goat anti-rabbit Alexa Fluor 647 (Thermo Fisher Scientific A21245; RRID: AB_2535813) at 1:500 was performed as previously described in the immunostaining section, to label neuronal bodies. Following the secondary staining of neurons, cell death in the injury core was then detected by terminal deoxynucleotidyl transferase dUTP nick end labeling (TUNEL) using the *in situ* cell death detection kit TMR red (Roche 12156792910) following manufacturer’s instructions. Brain sections were first permeabilized with cold PBS 1X, 0.1% Triton X-100, 0.1% sodium citrate for 2 min at 4°C. Sections were washed twice with PBS 1X and then incubated with TUNEL mix (label solution + enzyme terminal transferase) for 60 min at 37°C in a humidified chamber. Samples were washed 3 times with PBS 1X and mounted with SlowFade gold antifade with DAPI (Thermo Fisher Scientific S36939) using coverslips 22 mm × 50 mm 1.5 thickness and sealed with nail polish. Images were acquired in a Leica Thunder Imager 3D Tissue using a 20x/0.55 NA objective as 16-bit tile images at 7577 × 7577 pixels per tile (pixel size 0.325 × 0.325 μm) as 4.55 μm z-stacks (4 steps, 1.52 µm step size, with 10% overlap). The z-stacks were analyzed as orthogonal projections of maximum intensity with ImageJ software (RRID:SCR_003070). ROIs were selected by tracing the whole core of the injury using polygon tracing tool. Thresholds for the NeuN, TUNEL, and DAPI channels were determined using negative control sections and applied uniformly to all images. Colocalization between NeuN and TUNEL, and between DAPI and TUNEL, was assessed using the Image Calculator function in ImageJ. Results are presented as the TUNEL+ area colocalized with DAPI or NeuN, over the total DAPI or NeuN area, respectively. Representative images are 1500 × 1200 µm crops of maximum-intensity projections in the injury core.

### Magnetic Resonance Imaging (MRI)

Mouse MRI was performed to quantify lesion volume at 2 h, 1 day, 8 days, and 30 days following surgery. Axial two-dimensional multi-slice T2-weighted images were acquired using an M7 Compact MRI system (Aspect Imaging) with the following parameters: voxel size, 0.14 X 0.14 X 1 mm; 21 slices; no interslice gap; repetition time, 4300 ms; echo time, 50.45 ms; and vertical field of view, 20 mm. During imaging, mice were anesthetized with 1.5 – 2% isoflurane in oxygen and positioned in a dedicated mouse head holder (L25D23). Core body temperature was maintained at 37°C, and respiration was continuously monitored and maintained between 35 and 50 breaths per minute. Lesion volumes (mm³) were quantified using HOROS software (v4.0; RRID: SCR_017340).

### HEK cell transfection of DNA plasmids

HEK293T cells (ATCC CRL-3216; RRID:CVCL_0063) were plated on 10cm plates, coated with 0.1 mg/ml Poly-D-Lysine (Thermo Fisher Scientific A38904-01). Cells were maintained in DMEM (Thermo Fisher Scientific 11965092), Penicilin-streptomycin 10,000 U/mL (Thermo Fisher Scientific 15070063), GlutaMAX (Thermo Fisher Scientific 35050061) media at 37°C, 5% CO2 and ∼80% confluency. Plasmids AAV2- GfaABC1D-eGFP-4x6t and AAV2-GfaABC1D-Chrdl1-HA-4x6t were transfected in HEK293T cells using Lipofectamine 3000 reagent (Thermo Fisher Scientific L3000001) and OptiMEM (Thermo Fisher Scientific 31985-070), following manufacturers instructions. Each plasmid was transfected at 25 µg and cells were later collected 3 days after transfection for mRNA extraction and qPCR analysis to confirm expression of eGFP and Chrdl1-HA before use on tissue (Figure S3B).

### RNA extraction and real time quantitative PCR (RT-qPCR)

RNA was purified using the RNeasy Mini Kit according to the manufacturer’s instructions (Qiagen 74104). Primary neurons or HEK293T cultures were lysed in RLT buffer supplemented with 1% β-mercaptoethanol using cell scrapers (Chemglass Life Sciences CGN-2403), whereas adult mouse cortex and hippocampus were homogenized in the same buffer using a pellet pestle (DWK Life Sciences K749521-1590) and cordless motor (DWK Life Sciences K749540-0000). Lysates were passed through QIAshredder columns (Qiagen 79656), and RNA was isolated according to the manufacturer’s instructions. cDNA was generated using 200 ng of extracted mRNA and Superscript VILO MasterMix (Thermo Fisher Scientific 11755050), following manufacturer’s instructions and processed using a thermocycler (Bio Rad 1841100). The cDNA generated was used to perform qPCR. SYBR Green Master Mix (Thermo Fisher Scientific 4309155) was used for amplification and detection of Chrdl1, Syt1, GFAP, CSPG4, FgFr4, Iba1, and GAPDH (for primer sequences see Key Resources Table). Amplification and melting curves were measured with the Applied Biosystems 7500 real time PCR System (A&B 4351107). Cycle of quantification (Cq) values were thresholded by the A&B software and normalized to the average of the reference gene Gapdh. Normalized Cq values were then used to determine the fold change relative to control groups.

### Oxygen and glucose deprivation

Primary hippocampal neurons were subjected to oxygen and glucose deprivation (OGD) or normoxic control conditions for 20 min at DIV16. For OGD, cultures were transferred to glucose-free growth medium (Thermo Fisher Scientific, A1443001) and incubated at 37°C in a hypoxic chamber (Stem Cell technologies 27310) containing 95% N₂ and 5% CO₂. For normoxic controls, cultures were transferred to glucose-containing growth medium (Thermo Fisher Scientific, 21063029) and maintained at 37°C in a humidified incubator with 5% CO₂. Following the 20 min incubation, coverslips were either collected immediately or returned to their original growth medium and allowed to recover under normoxic conditions (37°C, 5% CO₂) for 72 h (referred to as reperfusion) before fixation and further procesing.

### Immunocytochemistry against cleaved caspase 3 (CC3) and analysis

Coverslips of primary hippocampal neurons were washed three times with PBS 1X and fixed with 4% PFA for 10 min, followed by three washes with PBS 1X. Cells were permeabilized with three washes of PBS 1X, 0.1% Triton X-100 for 5 minutes at room temperature, and blocked in 5% NGS with 0.1% Triton X-100 in PBS 1X for 1 h at room temperature. Coverslips were incubated overnight at 4°C with anti–CC3 at 1:400 (Cell Signaling Technology 9661; RRID:AB_2341188) in blocking buffer, washed three times with PBS 1X, 0.1% Triton X-100, and incubated with secondary antibody goat anti-rabbit Alexa Fluor 594 at 1:500 (Thermo Fisher Scientific A11037; RRID:AB_2534095) for 1 h at room temperature. After three additional PBS 1X, 0.1% Triton X-100 three washes, coverslips were mounted using SlowFade Gold antifade mountant with DAPI (Thermo Fisher Scientific, S36939) on SuperFrost Plus slides (VWR) and sealed with clear nail polish. Three to four ROIs were acquired per coverslip (three coverslips per condition) using a Leica Thunder Imager 3D Tissue microscope with a 20x/0.55 NA objective. Images were acquired as 16- bit images (2048 × 2048 pixels; pixel size 0.325 × 0.325 μm) as 7.58 μm z-stacks of 6 steps with 10% overlap. Quantification of cell death was performed blinded to treatment and condition using ImageJ software (RRID: SCR_003070). A single fluorescence intensity threshold was established by manually adjusting the threshold on representative images in ImageJ and was applied uniformly across all images to identify DAPI+ nuclei and CC3+ cells. Cell death was calculated as the % of CC3+ cells relative to the total number of DAPI+ nuclei within each ROI.

### Surface and intracellular GluA2 staining on primary neurons and analysis

For endogenous surface and intracellular staining of GluA2, after Chrdl1 or vehicle (0.1% BSA 4 mM HCl) treatment, primary hippocampal cell cultures were first fixed for 8 min with 4% PFA. Surface GluA2 was labeled with primary antibody against the mouse N-terminal extracellular domain of GluA2 (Millipore MAB397; RRID:AB_2113875) at 1:50 dilution in antibody buffer. Coverslips were then washed three times with PBS 1X for 5 min at room temperature. Coverslips were incubated with secondary antibody goat anti-mouse Alexa 488 (Thermo Fisher Scientific A11029; RRID:AB_2534088) at 1:500 dilution in antibody buffer. This was followed by a permeabilization step using 0.2% Triton X-100 in 10% goat serum in 150mM NaCl (Sigma-Aldrich S9625), 50mM Tris (Sigma-Aldrich 252859), 100mM L-lysine (Thermo Fisher Scientific 56871), and 1% BSA (Sigma-Aldrich A4161) for 1 h at room temperature. After permeabilization, coverslips were incubated with a fresh 1:50 dilution of the same anti-GluA2, followed by incubation with secondary antibody goat anti-mouse Alexa 647 (Thermo Fisher Scientific A21235; RRID:AB_2535804) at 1:500 dilution to label intracellular GluA2. After three washes with PBS 1X, coverslips were mounted using SlowFade Gold antifade mountant with DAPI (Thermo Fisher Scientific, S36939) on SuperFrost Plus slides (VWR) and sealed with clear nail polish. Images were acquired on a Leica SP5 confocal microscope (RRID:SCR_020233) using a 63x/NA 1.4 oil-immersion objective as 8-bit images at 1024 x 1024 pixels (pixel size 120.1 x 120.1 nm) as a 3.78 µm-thick z-stack of 11 steps with 0.38 µm step size, and 10% overlap. Images were analyzed using ImageJ/Fiji (RRID:SCR_003070). There were three coverslips for each experimental condition, with three neurons imaged per coverslip. Values from 3-4 randomly selected 25-µm sections of secondary dendrites were averaged to obtain a single value per neuron. For each independent neuronal culture, measurements from OGD-treated groups were normalized to their corresponding control group to minimize variability between neuronal cultures. Images were thresholded to identify GluA2+ clusters, and a minimum particle size threshold of 0.12 μm^2^ was applied using the Analyze Particles function to exclude background signal. Surface and intracellular GluA2 were quantified as the number of surface or intracellular GluA2+ puncta divided by the total number of GluA2 puncta (surface + intracellular) within each dendritic segment.

### Behavioral testing

All behavioral testing was conducted with the experimenter blinded to condition, and mice were tested in randomized order. Three mice (one eGFP and two Chrdl1 OE) that did not survive to the final experimental time point were excluded from all analyses. Prior to behavioral testing, mice were handled for 3 – 5 min per day for 2 consecutive days. Before all behavioral assays, mice were acclimated to the testing room and lighting conditions for at least 30 min. For baseline experiments, behavioral testing began 2 weeks after retro-orbital AAV injection and prior to surgery. Lighting and temperature (20 – 24°C) were controlled in the testing room, and insulated from external noise. All sessions were recorded using a Logitech BRIO camera (1080p–4K resolution), and animal position was tracked using ANY-maze software (v7.48). Mice that died during the study (Figure S3I) were excluded from statistical analyses.

#### Open field

Open field was used to evaluate exploratory activity, locomotion, and anxiety-like behavior^36^. Testing was performed in a square open field arena (45 × 45 × 45 cm) located in the dedicated behavioral testing room under bright lighting with no visual cues. Mice were handled daily for 2–3 consecutive days prior to testing and acclimated to the testing room for 30 min before each session. Two days before testing, mice were habituated to the arena by free exploration for 3 min per day, with no data collected during habituation. On the testing day, mice were again acclimated to the room for 30 min and then placed in the center of the arena to freely explore for 5 min. Animal movement was recorded using an overhead camera and analyzed using ANY-maze software (v7.48). The arena was cleaned with Peroxigard between animals to eliminate olfactory cues. The arena was divided into a grid (7.5 × 7.5 cm squares), defining a central zone (16 squares) and a peripheral zone (20 squares). Quantified parameters included total distance traveled (m), % time spent in center, immobile time (s, not moving for more than 2000 ms) and average speed (m/s). Baseline measurements were acquired 8 days prior to stroke and 1,8, and 30 dps.

#### Novel object recognition test (NORT)

Novel object recognition test (NORT) was performed in the same arena (45 × 45 × 45 cm), with the arena position maintained constant throughout testing. Mice were handled for 2 days prior to testing and acclimated to the testing room for at least 30 min under dim indirect lighting conditions. During the familiarization phase, two identical objects were placed equidistant from the arena walls, and mice were allowed to explore freely for 10 min. After a 15 –18 min retention interval, one familiar object (FO) was replaced with a novel object (NO) differing in shape, color, and texture, and mice were reintroduced into the arena and allowed to explore for 5 min. Animal position was tracked using ANY-maze software (v7.48). Exploratory behavior was defined as nose-directed investigation at a distance ≤2 cm from the object. Time spent and distance traveled exploring each object were quantified. The discrimination index was calculated as (NO exploration time − FO exploration time) divided by total exploration time during the test session. Recognition index was calculated as NO exploration time divided by total exploration time. Measurements were obtained on the same day as open field testing, following re-acclimation under dim, indirect lighting

#### Barnes maze

Spatial learning and memory were assessed using the Barnes maze^37^. The maze consisted of a white circular platform (92.7 cm diameter) with 20 evenly spaced circular holes (4.5 cm diameter) around the perimeter, elevated approximately 114 cm above the floor. One hole led to a white opaque escape box (23 × 10 × 6 cm). Visual cues were positioned on the walls approximately 30 cm from the platform and remained constant throughout testing. Bright lighting was used to motivate escape behavior. Mouse position was tracked using ANY-maze software (v7.48). Mice were acclimated to the escape box for 3 minutes prior to testing to familiarize them with it. Mice underwent three 3-min training trials per day for four consecutive days, with an intertrial interval of approximately 1 h. Mice that failed to enter the escape box within 3 min were gently guided to it. Training performance was quantified using path efficiency, path length to locate the target box, latency to first locate the target box, and number of errors. A probe trial was conducted 24 h after the final training session, during which the escape box was removed, and mice were allowed to explore freely for 180 s. Each training and probe testing session (baseline [location A], subacute phase [location B], chronic phase [location C]) was completed using a different box location in a different quadrant of the platform. Latency and path length to the target box, as well as prior target box locations, were quantified. Path efficiency was calculated as the direct path distance to box (0.3 m) divided by the path length to first investigate box. Number of errors was defined as the number of times that the mouse investigated other locations prior to investigating the target box.

### Synaptosome and cytosolic fraction preparation

Synaptosome and cytosolic fractions were prepared from WT or Chrdl1 global KO^14^ male mouse cortex or hippocampus using differential centrifugation. Freshly dissected tissue was snap-frozen and homogenized on ice in Syn-PER™ Synaptic Protein Extraction Reagent (Thermo Fisher Scientific 87793) supplemented with protease inhibitors (Roche 04693116001), 1 tablet per 10 mL, at a ratio of 10 mL buffer per gram of tissue. Tissues were dounce-homogenized with ∼10 slow strokes and centrifuged at 1,200 x g for 10 min at 4 °C to remove nuclei and debris. The supernatant was transferred to a new tube and centrifuged at 15,000 x g for 20 min at 4 °C to pellet synaptosomes. The resulting supernatant was retained as the cytosolic fraction, while the synaptosome pellet was resuspended in Syn-PER reagent. For proteomic analyses, synaptosome pellets were solubilized in sodium deoxycholate (Na-DOC; 0.2% final) (Sigma-Aldrich D6750), rotated end-over-end for 1 h at 4°C, sonicated on ice (20% amplitude, 2–3 pulses of 8–10 s), and clarified by centrifugation at 15,000 x g for 5 min at 4°C. Supernatants were then collected for protein quantification and mass spectrometry.

### Western blot

Samples for western blot were prepared as 20 µg of total protein mixed with 1X LDS sample buffer (Thermo Fisher Scientific B0007) and reducing agent (Thermo Fisher Scientific B0009), denatured at 95°C for 5 min, and separated on 4 – 12% Bis-Tris polyacrylamide gels (Thermo Fisher Scientific NW04120BOX) using MES running buffer (Thermo Fisher Scientific B0002). We use the PageRuler™ Plus Prestained Protein Ladder (10 – 250 kDa; Thermo Fisher Scientific 26619) as a molecular weight standard. Electrophoresis was performed at 150 V until adequate protein separation was achieved. Proteins were transferred onto Immobilion®-P PVDF membranes (Sigma Millipore IPVH00010) using a wet transfer system in Bolt transfer buffer (Thermo Fisher Scientific BT00061) with 10% methanol and antioxidant (Thermo Fisher Scientific BT0005) at 100 V for 2 h. Membranes were rinsed in Tris-buffered saline (TBS) (Sigma T5912), blocked for 1 h at room temperature in casein-based blocking buffer (Thermo Fisher Scientific 37583), and incubated overnight at 4°C with primary antibodies mouse anti-GluA2 (Millipore MAB397) at 1:1,000 or mouse anti-GAPDH (Proteintech 60004-1-lg) at 1:10,000 dilution in TBS + 0.1% Tween-20 (Thermo Fisher Scientific J20605-AP) (TBS-T). Following three washes in TBS-T, membranes were incubated with secondary antibody goat anti-mouse Alexa Fluor 680 (Invitrogen A32729) at 1:10,000 for 2 h at room temperature. Blots were washed again and imaged using an Odyssey Li-Cor infrared imaging system (LI-COR Biosciences 9142).

### Liquid chromatography tandem mass spectrometry (LC-MS/MS)

Cytosolic or synaptosome samples (20 µg) were sent to the Wistar Institute to perform liquid chromatography tandem mass spectrometry. Samples were run 0.5 cm into a NuPAGE 10% Bis-Tris gel (ThermoFisher Scientific NP0301BOX) and lanes were excised and subjected to in-gel trypsin digestion. Digests were resuspended in 0.015% dodecyl maltoside, 0.1% trifluoroacetic acid before analysis. LC-MS/MS analysis was performed using a Vanquish Neo UHPLC system coupled to an Orbitrap Astral mass spectrometer (ThermoFisher Scientific BRE725660). Samples were injected onto an Acclaim Pepmap C18 trap column (100 Å, 75 μm i.d. x 2 cm packed with 3 μm C18 resin, Thermo Fisher Scientific 164946) and separated on a NanoEase Symmetry BEH C18 nanocapillary analytical column (130 Å, 75 μm i.d. x 25 cm, 1.7 μm particle size; Waters 186008795). Buffer A consisted of 0.1% formic acid in Milli-Q water, and buffer B consisted of 0.1% formic acid in acetonitrile. Separation was achieved using a 40 min linear gradient 5- 30% B over 25 min, 30-40% B over 6 min, 40-80% B over 2 min, followed by a 7 min hold at 80% B before re-equilibration.

MS data were acquired in data-independent acquisition mode. An MS1 scan was collected every 0.6 s in the Orbitrap at 240,000 resolution. Ions were injected for 3 ms or until an AGC target of 5e6 was reached. Precursor ions within a mass range of 380-980 m/z were collected in 2 m/z isolation windows with a maximum injection time of 3 ms or until an AGC target of 5e4 ions was reached. Precursors were fragmented using 25% normalized collisional energy. MS2 scans from 110 – 2000 m/z were collected in the Astral mass analyzer.

## QUANTIFICATION AND STATISTICAL ANALYSIS

Sample sizes were determined a priori by power analysis (α = 0.05, power = 0.8) to ensure adequate statistical power to detect biologically relevant effects. Statistical analyses and figure generation were performed using GraphPad Prism v10 (RRID:SCR_002798). We used unpaired two-tailed Student’s t-tests for comparisons between two groups, one-way ANOVA for comparisons among more than two groups, and two-way ANOVA for experiments involving two independent variables. When multiple comparisons were performed, Holm–Šídák or Fisher’s least significant difference (LSD) post hoc tests were used, as appropriate. Statistical tests, sample sizes, and significance values for individual experiments are reported in the corresponding figure legends.

### Proteomic Data Analysis

MS RAW files were searched with DIA-NN (v2.2.0; RRID:SCR_022865)^38^ against a spectral library generated from the UniProt mouse database with curated isoforms (UP000000589, downloaded 4/25/2025; RRID:SCR_002380) supplemented with a contaminant database. N-terminal acetylation, N-terminal methionine excision, and methionine oxidation were set as variable modifications. Cysteine carbamidomethylation was set as a static modification. Data were searched with full tryptic specificity allowing for 2 missed cleavages, 1 variable modification, a peptide length of 7 – 50 amino acids, a precursor charge of 1 – 4, and a scan window of 15. MS1 and MS2 mass accuracy were set to 5.0 and 10.0 ppm, respectively. The match-between-runs feature and the RT-dependent cross-run normalization feature were both enabled. Precursor, peptide, and protein false discovery rates were controlled at 1%. Contaminants, proteins identified by a single peptide, and proteins without at least 3 valid values in at least one of the eight experimental groups were removed, yielding a filtered dataset of 8,948 protein groups across the 24 samples. Non-normalized abundances and per-sample stripped-peptide counts were carried forward.

Downstream analysis was performed via a Nextflow (v25.10.4; RRID:SCR_024135)^39^ pipeline implemented in Python (v3.12.13; RRID:SCR_008394) and R (v4.5.3; RRID:SCR_001905). Each pairwise comparison was analyzed independently, starting from this filtered dataset. Within each comparison, protein groups detected in at least 2 of the 3 replicates of at least one of the two groups being compared were retained, log2 transformed, and median normalized (8,339–8,666 protein groups per comparison). Missing values were classified as missing-not-at-random or missing-at-random and imputed accordingly: values in a group falling below the detection threshold (>1 of 3 replicates missing) were treated as left-censored and imputed by MinProb (0.01 quantile, width 0.3 SD), and remaining sporadic missing values were imputed by distance-weighted k-nearest neighbors over proteins (k = 10) using scikit-learn (v1.8.0; RRID:SCR_002577)^40^.

Sample relationships across the full dataset were visualized by principal component analysis. Because filtering, imputation, and normalization were performed independently within each pairwise comparison, the overview PCA was computed instead on the quantile-normalized, minimum-value-imputed matrix provided by the proteomics core, comprising all 8,948 filtered protein groups across the 24 samples. The 500 protein groups with the highest variance across samples were retained, and principal components were computed on protein-centered abundances without scaling (scikit-learn). Principal components 1 and 2 are shown. This matrix was used for this analysis only; all differential abundance and enrichment analyses used the median-normalized, MinProb/kNN-imputed matrices described above.

Differential abundance was assessed with limma (v3.66.0; RRID:SCR_010943)^41^ followed by DEqMS (v1.28.0; RRID:SCR_025605)^42^, which moderates variance by peptide count, using empirical Bayes. In both hippocampal comparisons, one outlier wild-type replicate (HIPsynap_WT-3 and HIPcyto_WT-3, respectively) was downweighted in the primary analysis using a mean-shift indicator in the design matrix. P-values were adjusted using Benjamini-Hochberg correction, and proteins with an absolute fold change ≥ 2 and an adjusted p-value < 0.05 were considered significant.

Preranked gene set enrichment analysis was performed with GSEApy (v1.1.13; RRID:SCR_025803)^43^ against the mouse MSigDB GO biological process collection (v2026.1.Mm; RRID:SCR_016863), using gene sets of 15-500 genes, the DEqMS moderated t-statistic as the ranking metric, and 1,000 permutations. Significance was assessed using the permutation-based false discovery rate, and terms with q < 0.05 were considered significant. Terms with q < 0.05 were considered significant; the ten such terms with the largest absolute normalized enrichment score per comparison are shown.

## ADDITIONAL RESOURCES

Readers may download a detailed protocol for the hippocampal photothrombosis from https://link.springer.com/protocol/10.1007/978-1-0716-2926-0_4. Please cite as Blanco-Suárez E. Photothrombotic Model to Create an Infarct in the Hippocampus. Methods Mol Biol. 2023;2616:29-38. doi: 10.1007/978-1-0716-2926-0_4. PMID: 36715925.

## RESOURCES TABLE

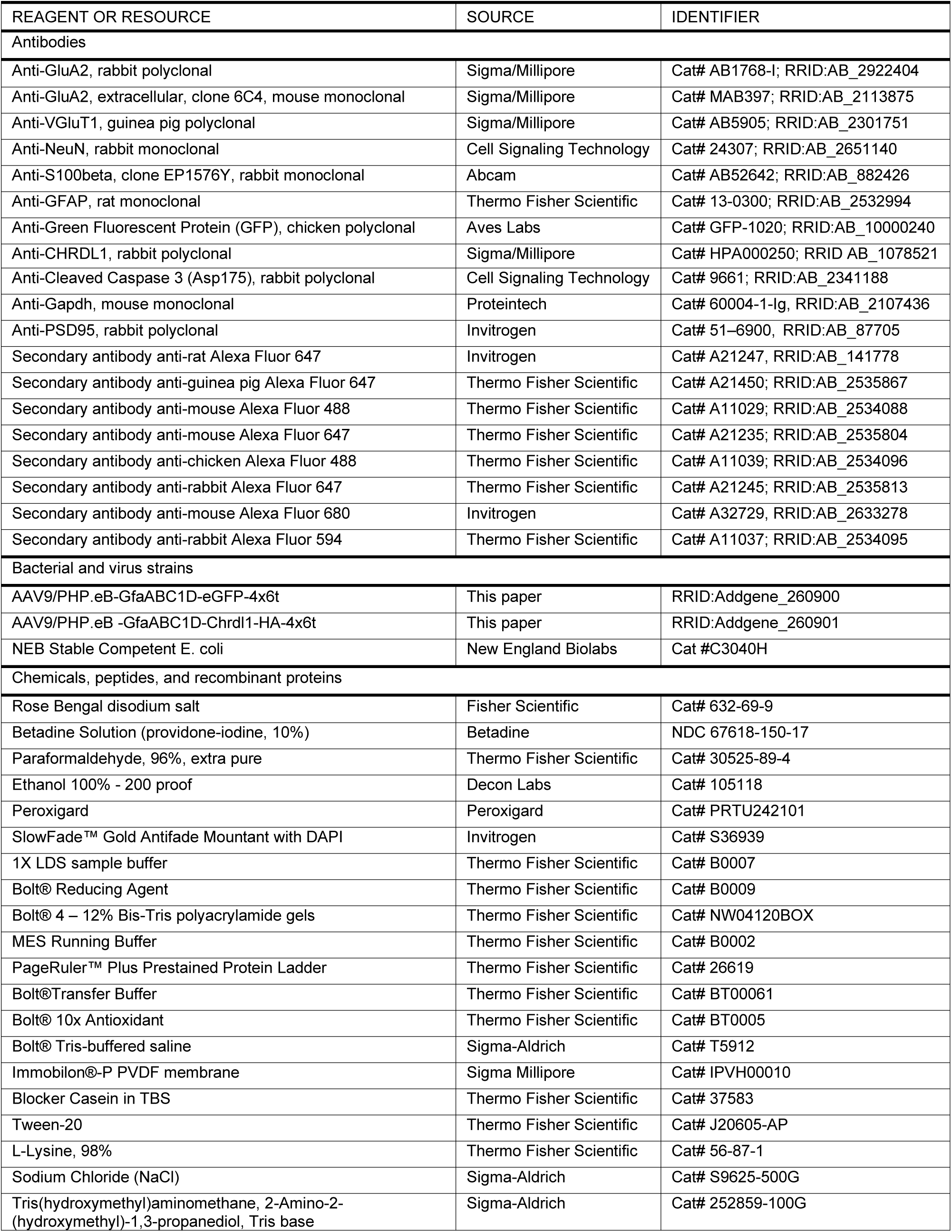
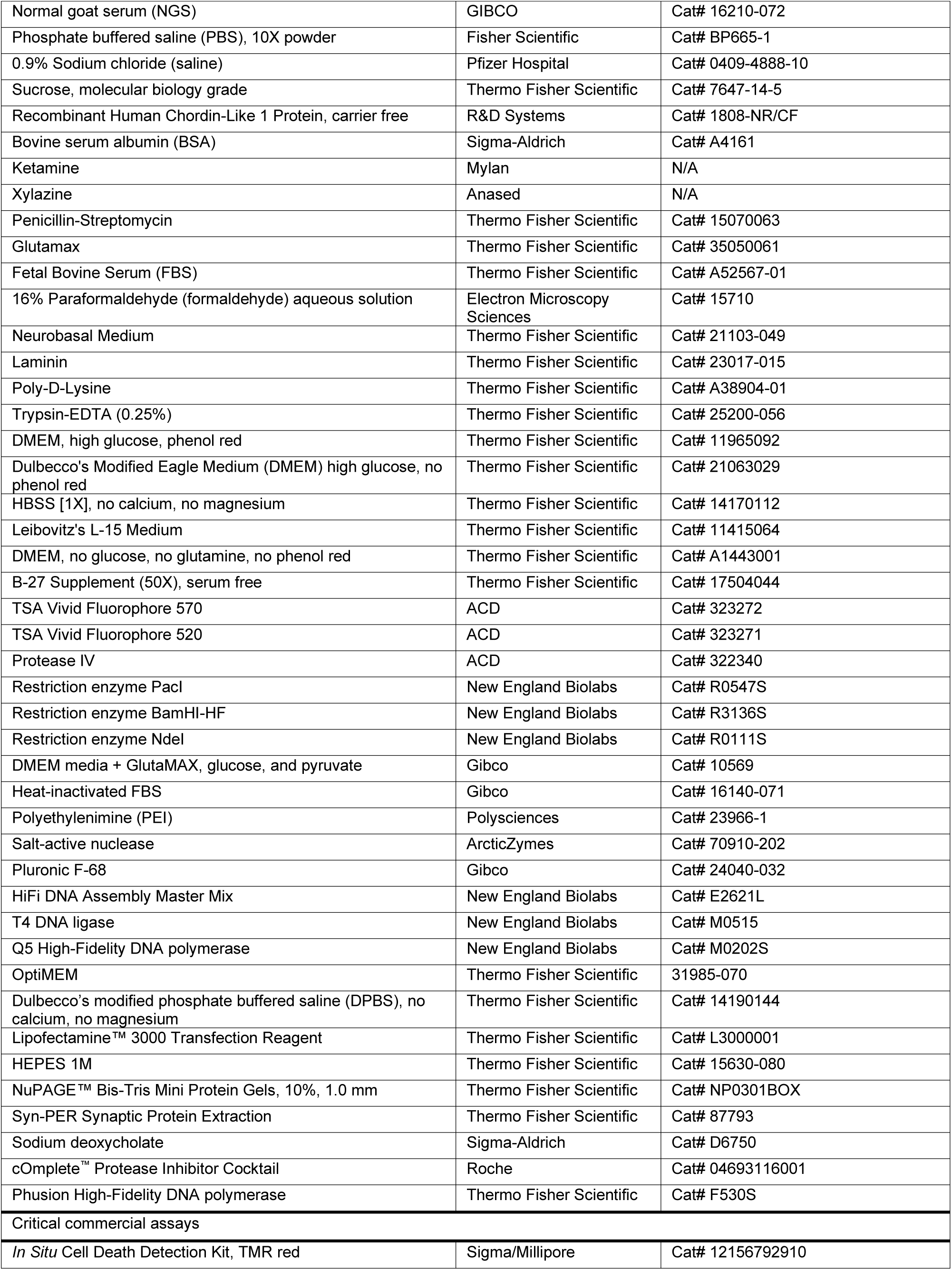
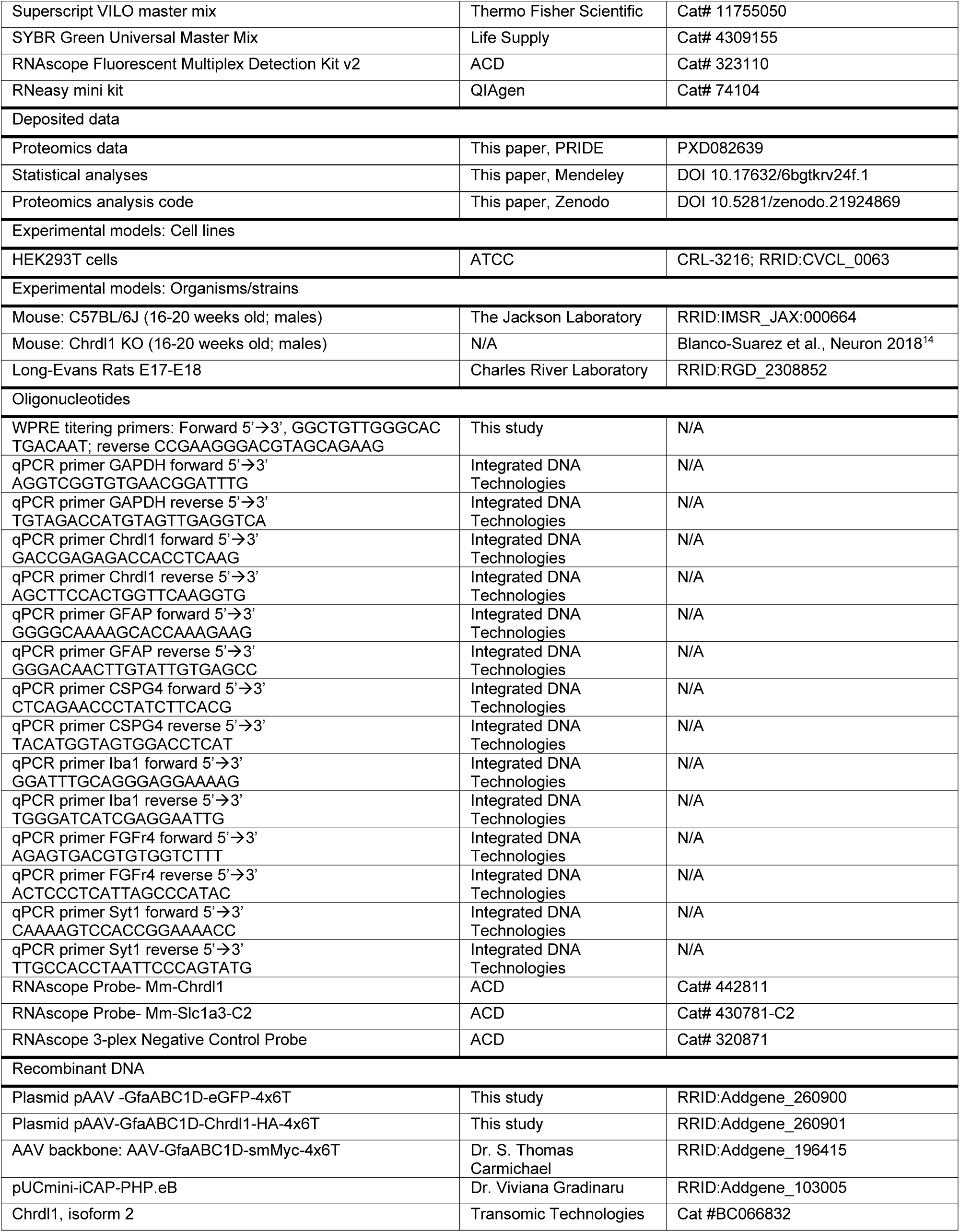
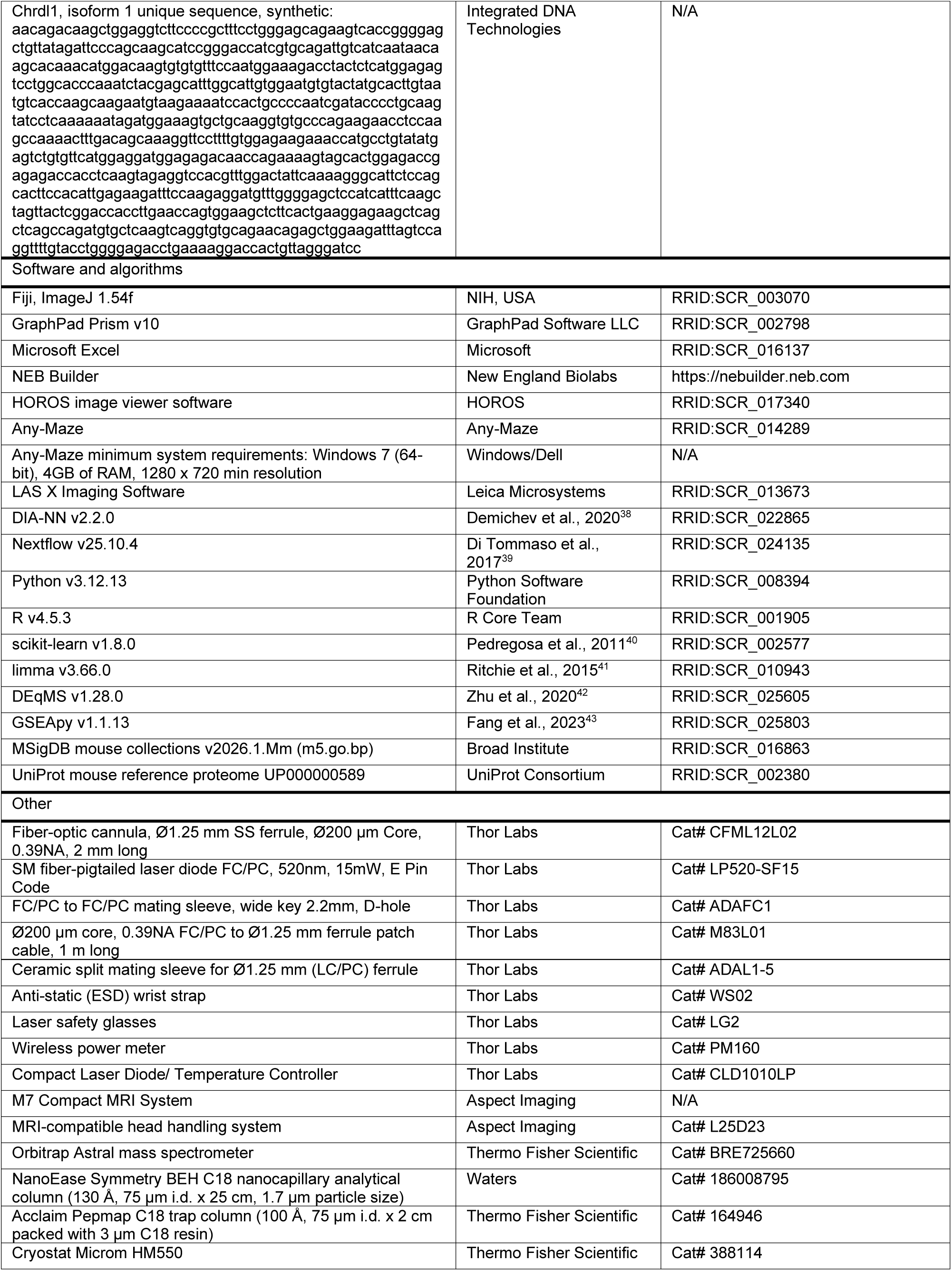
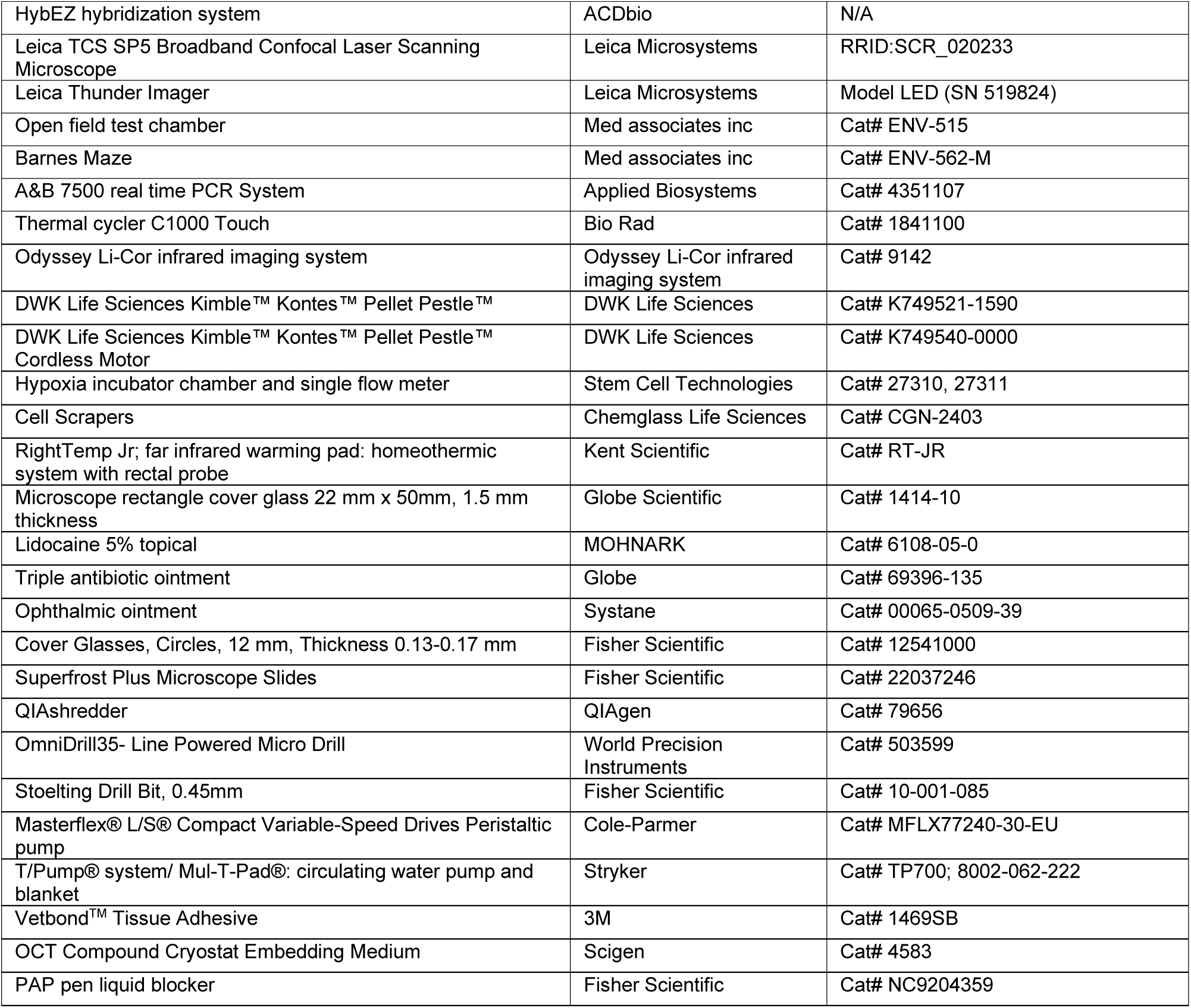

---

**Figure S1.**
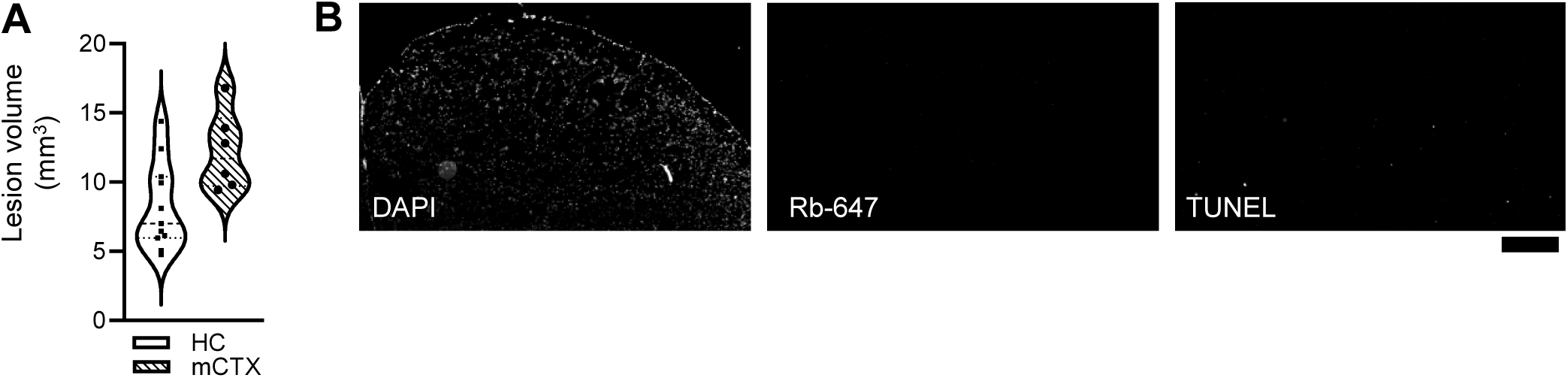
Photothrombotic lesion volume and staining controls. A) MRI-based quantification of lesion volume 2 h post-stroke (hps) following photothrombosis in the motor cortex (mCTX, n = 6) or hippocampus (HC, n = 11). B) Representative negative controls for TUNEL (enzyme omitted) and immunohistochemistry (primary antibody omitted) in the motor cortex 1 day post-stroke (dps). Scale bar 500 μm. Related to Figure 1.

**Figure S2.**
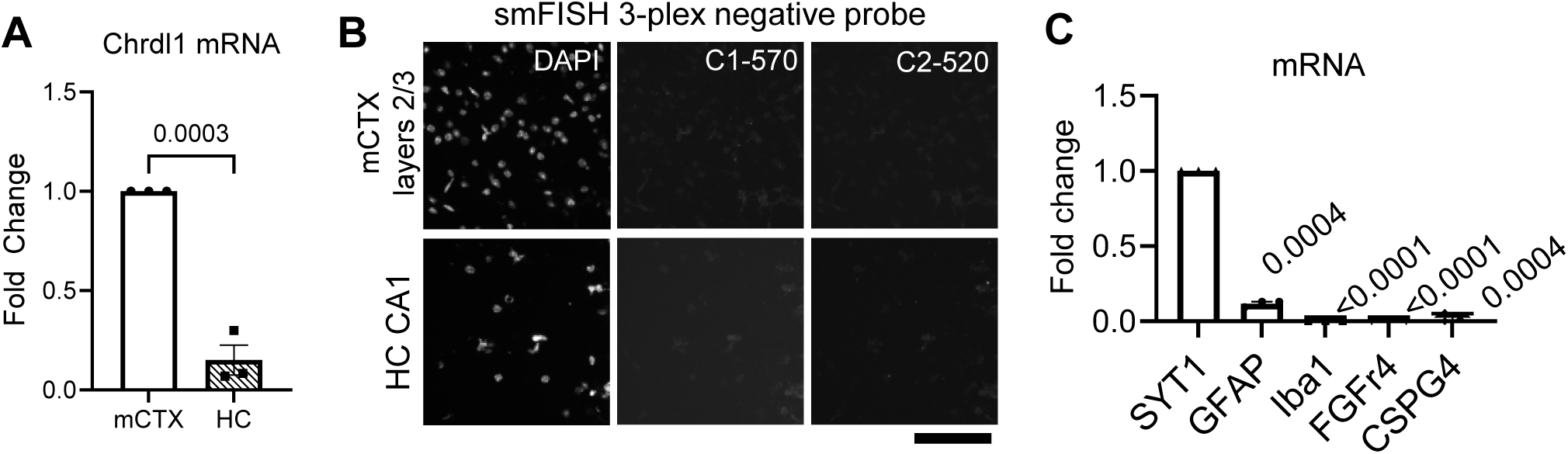
Validation of cell-type markers and RNA detection controls. A) RT-qPCR quantification of Chrdl1 mRNA in adult WT cortex (n = 3) and hippocampus (n = 3). Data represented as mean ± SEM with individual data points per mouse. Statistics by unpaired two-tailed t test. P-value on graph. B) Representative smFISH negative control probe in upper cortical layers 2/3 (top) and hippocampal CA1 (bottom). Scale bar 50 μm. C) RT-qPCR validation of primary hippocampal neuronal cultures (DIV18–21). Expression of neuronal (Syt1), astrocytic (Gfap), microglial (Iba1), fibroblast (Fgfr4), and oligodendrocyte precursor (Cspg4) markers relative to Gapdh (n = 3). Statistics by one-way ANOVA, followed by Holm- Šídák test, p-values on the graph. Related to Figure 2.

**Figure S3.**
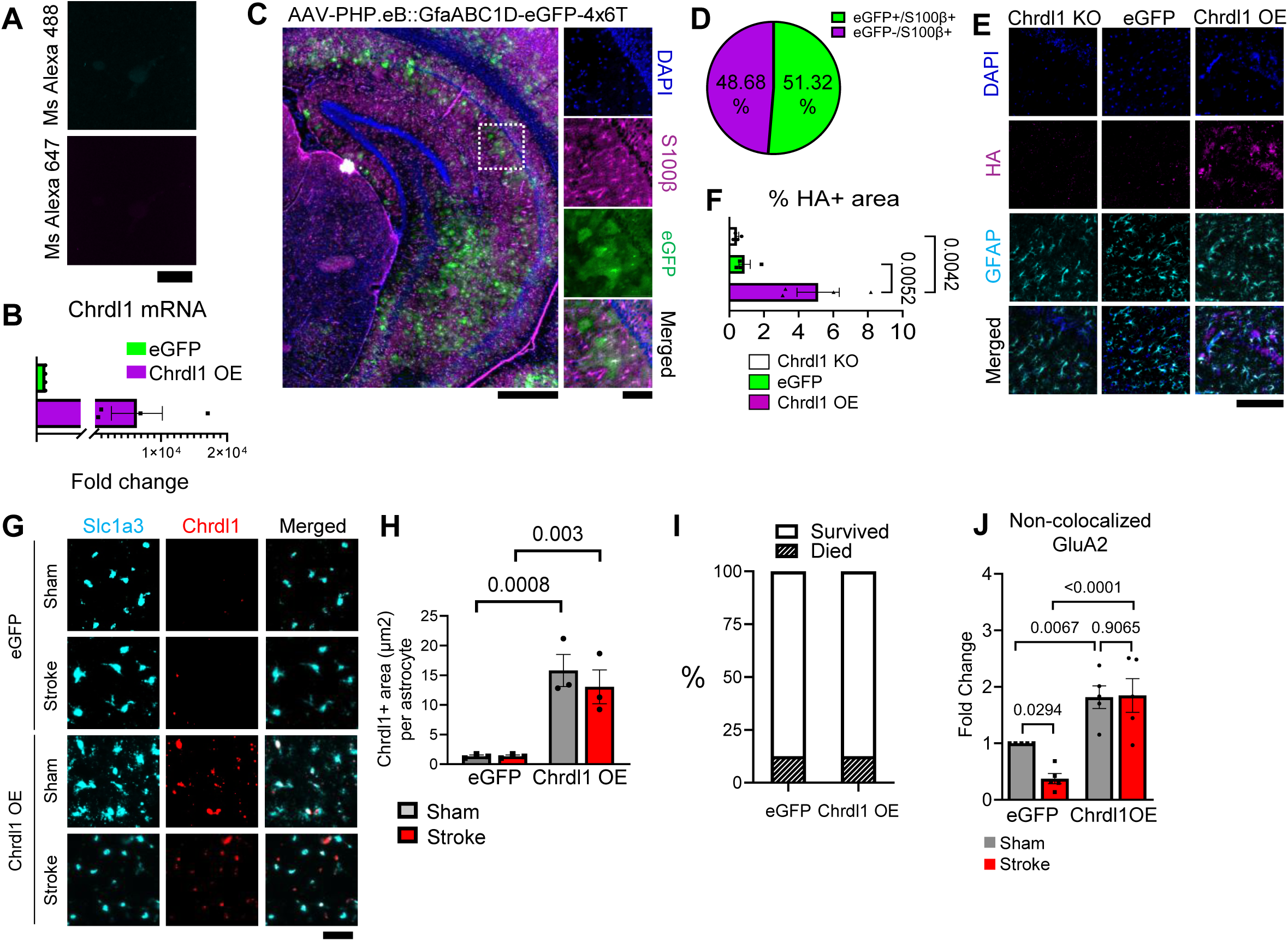
Validation of viral-mediated Chrdl1 overexpression and immunostaining. A) Representative negative controls for surface and permeabilized immunostaining in cultured hippocampal neurons. Primary antibody was omitted. Scale bar 25 μm. B) RT-qPCR validation of Chrdl1 overexpression in HEK293T cells transfected with AAV-eGFP or AAV-Chrdl1 OE plasmids (n = 4). C) Overview of a coronal section from an AAV-eGFP mouse. Scale bar 500 µm. Zoom-in images of dashed white box in CA1 area on the right, showing S100β (astrocyte marker), eGFP, DAPI, and all channels merged. Scale bar 100 µm. D) Quantification of S100β+ astrocytes expressing eGFP in the hippocampal CA1 following AAV- PHP.eB::GfaABC1D-eGFP-4x6T administration. E) Representative immunohistochemistry for HA (magenta) and GFAP (cyan) in the hippocampal CA1 of Chrdl1 KO mice injected with AAV-eGFP or AAV- Chrdl1 OE, 2 weeks after viral administration. Scale bar 200 μm. F) Quantification as % of HA+ area in the CA1 hippocampus ROIs (n = 4). G) Representative images of smFISH of Slc1a3 (astrocytes, cyan) and Chrdl1 (red) in the CA1 of the hippocampus from mice 2 weeks after AAV-eGFP (eGFP, AAV- PHP.eB::GfaABC1D-eGFP-4x6T) or AAV-Chrdl1OE (Chrdl1 OE, AAV-PHP.eB::GfaABC1D-Chrdl1-HA-4x6T) administration. Scale bar 50 µm. H) Quantification of G as Chrdl1+ area per Slc1a3+ cell in the CA1 hippocampus at 2 hps in the peri-infarct area of eGFP or Chrdl1 OE mice (N = 3 per condition). I) Survival following AAV administration and photothrombotic surgery. J) Quantification of extrasynaptic GluA2 puncta (non-colocalized with VGluT1), normalized to WT sham (n = 5). Bar graphs show mean ± SEM with individual data points representing biological replicates. Statistical analyses were performed using one-way ANOVA followed by Holm–Šídák multiple-comparisons test (F) or two-way ANOVA followed by uncorrected Fisher’s LSD multiple-comparisons test (H, J). Exact p-values are indicated on the graphs. Related to Figure 3.

**Figure S4.**
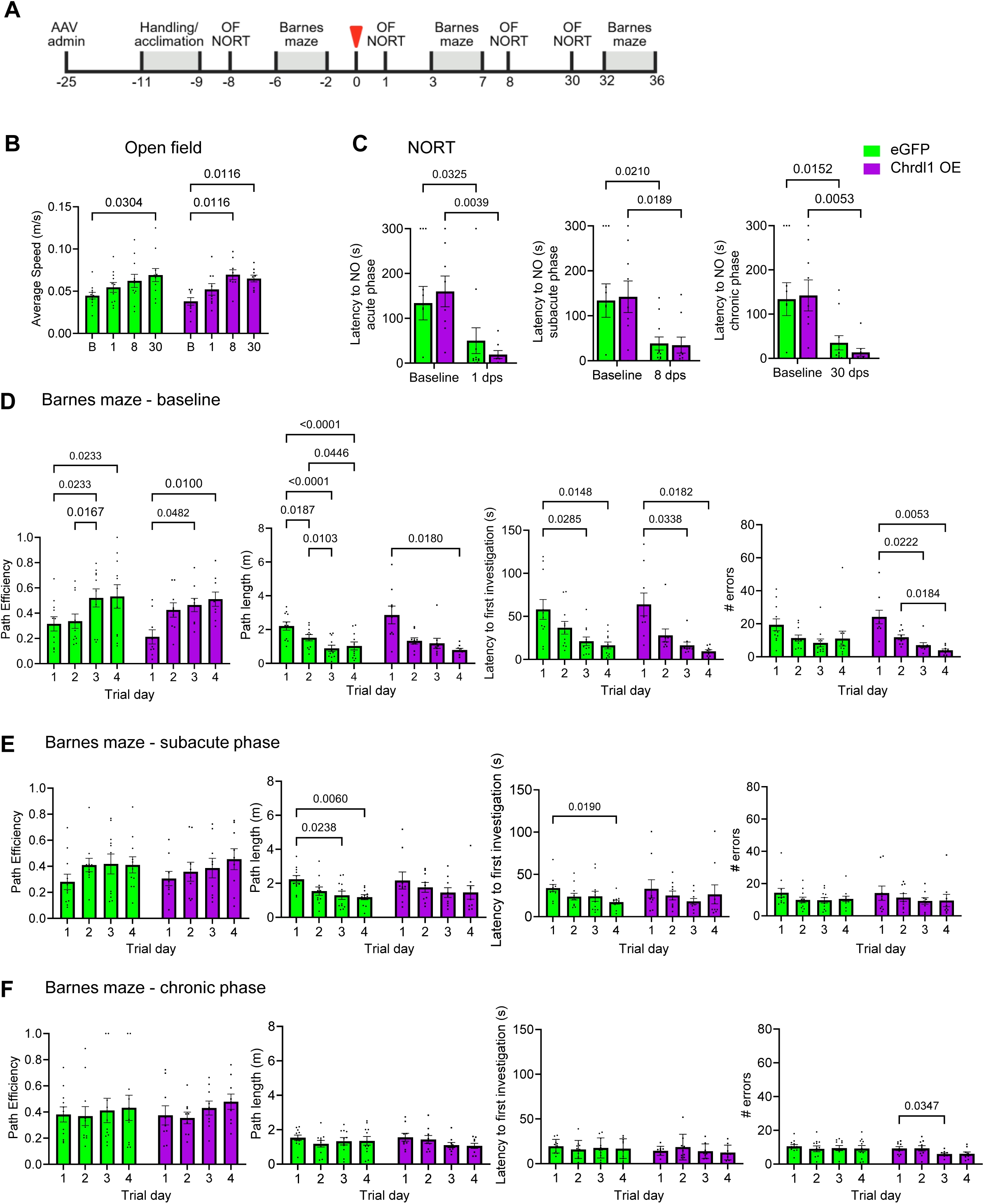
Overexpression of Chrdl1 does not alter baseline or post-stroke motor and cognitive behavioral outcomes. A) Experimental timeline for behavioral testing. The red marker indicates when mice underwent stroke surgery. B) Open field behavior test at baseline (B), 1 dps, 8 dps, and 30 dps, showing the quantified average speed (m/s) of eGFP (n = 11) and Chrdl1 OE (n = 9) mice, corresponding to experimental groups in Figures 5A – C. C) Novel object recognition test (NORT) in the acute (1 dps), subacute (8 dps), and chronic (30 dps) phases compared to baseline testing, showing the quantified latency to the novel object (s) for eGFP (n = 11) and Chrdl1 OE (n = 8 – 9) mice, corresponding to experimental groups in Figures 5D–5F. D–F) Barnes maze performance across four consecutive trial days at D) baseline (–6 to –3 days before stroke), E) subacute phase (3 to 6 dps), and F) chronic phase (32 to 35 dps) for eGFP (n = 11) and Chrdl1 OE (n = 9) mice. Quantified metrics of path efficiency, path length (m), latency (s), and errors committed before first locating the target hole, corresponding to experimental groups in Figures 5G– 5H. Bar graphs represent mean ± SEM with individual data points per mouse. Statistics by two-way ANOVA, followed by Holm-Šídák (B) or uncorrected Fisher’s LSD multiple-comparisons test (C – F) with only significant values (p < 0.05) indicated on the graphs for clarity.

**Figure S5.**
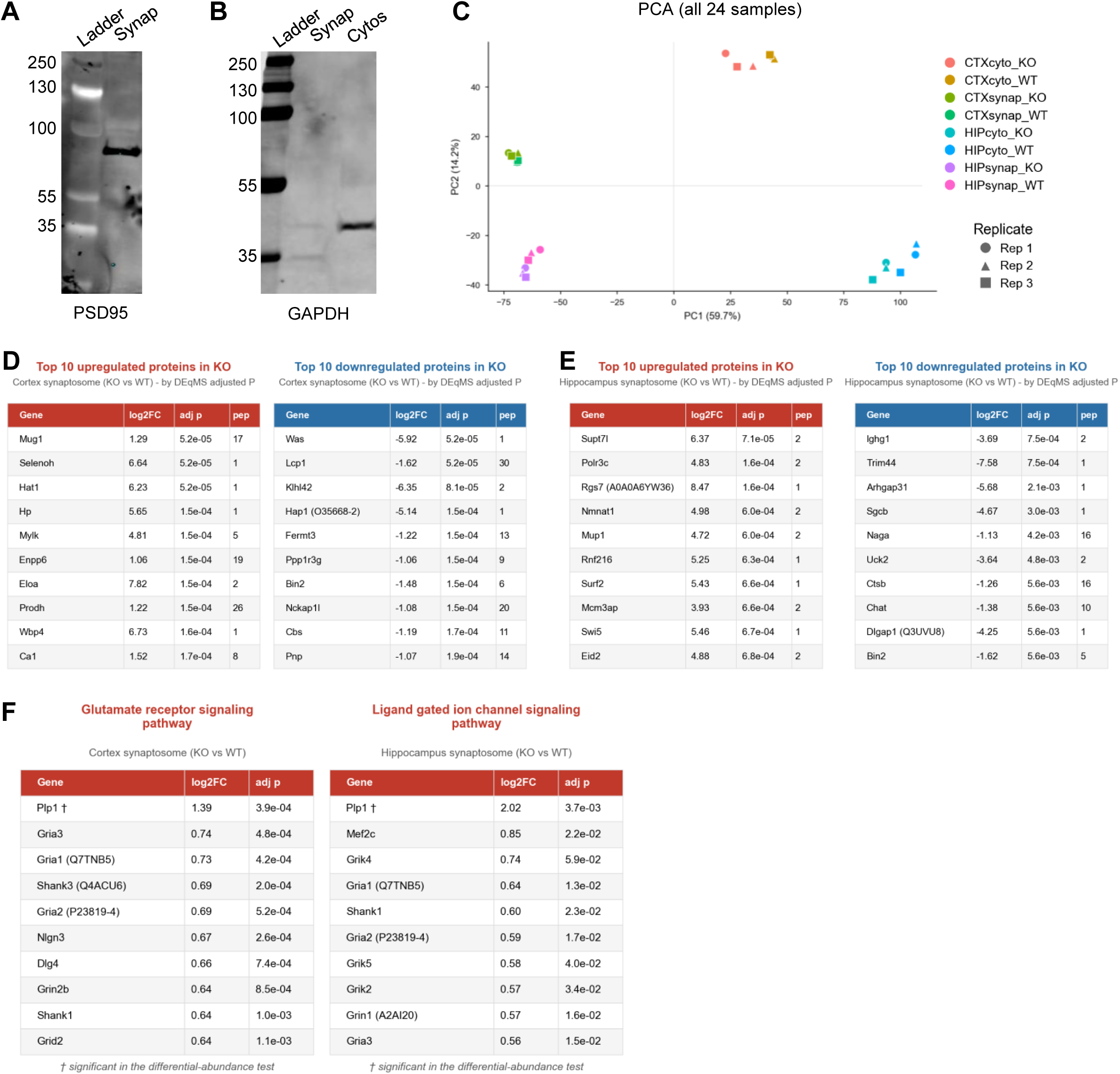
Validation of proteomic preparations and additional proteomic analyses. A – B) Representative immunoblots validating synaptosome (synap) and cytosolic (cytos) fractionation using PSD95 (A) and GAPDH (B) as synaptic and cytosolic markers, respectively. C) Principal component analysis of all proteomic samples based on the 500 most variable proteins. D – E) Top ten proteins upregulated and downregulated in Chrdl1 KO cortical (D) and hippocampal (E) synaptosomes ranked by adjusted DEqMS *P* value. F) Leading-edge proteins contributing to glutamatergic pathway enrichment in cortical synaptosomes and ligand-gated ion channel signaling in hippocampal synaptosomes. Related to Figure 6.

**Table S1.** Data points used in all statistical analyses in Figures 1 – 5, S1 – 4.

## REFERENCES

1 Cramer, S. C. & Chopp, M. Recovery recapitulates ontogeny. Trends in Neurosciences 23, 265– 271 (2000). 10.1016/S0166-2236(00)01562-9

2 Allen, N. J. Role of glia in developmental synapse formation. Current Opinion in Neurobiology 23, 1027–1033 (2013). 10.1016/j.conb.2013.06.004

3 Boyle, B. R., Berghella, A. P. & Blanco-Suarez, E. Astrocyte Regulation of Neuronal Function and Survival in Stroke Pathophysiology. Adv Neurobiol 39, 233–267 (2024). 10.1007/978-3-031-64839-7_10

4 Collyer, E. & Blanco-Suarez, E. Astrocytes in stroke-induced neurodegeneration: a timeline. Frontiers in Molecular Medicine 3 (2023).

5 Gleichman, A. J. & Carmichael, S. T. Astrocytic therapies for neuronal repair in stroke. Neuroscience Letters 565, 47–52 (2014). 10.1016/j.neulet.2013.10.055

6 Blanco-Suarez, E. & Hanley, J. G. Distinct Subunit-specific α-Amino-3-hydroxy-5-methyl-4- isoxazolepropionic Acid (AMPA) Receptor Trafficking Mechanisms in Cultured Cortical and Hippocampal Neurons in Response to Oxygen and Glucose Deprivation. Journal of Biological Chemistry 289, 4644–4651 (2014). 10.1074/jbc.M113.533182

7 Dixon, R. M., Mellor, J. R. & Hanley, J. G. PICK1-mediated Glutamate Receptor Subunit 2 (GluR2) Trafficking Contributes to Cell Death in Oxygen/Glucose-deprived Hippocampal Neurons. Journal of Biological Chemistry 284, 14230–14235 (2009). 10.1074/jbc.M901203200

8 Guo, C. & Ma, Y.-Y. Calcium Permeable-AMPA Receptors and Excitotoxicity in Neurological Disorders. Frontiers in Neural Circuits 15 (2021).

9 Liu, B. et al. Ischemic insults direct glutamate receptor subunit 2-lacking AMPA receptors to synaptic sites. J Neurosci 26, 5309–5319 (2006). 10.1523/jneurosci.0567-06.2006

10 Le, D. A. et al. Caspase activation and neuroprotection in caspase-3- deficient mice after in vivo cerebral ischemia and in vitro oxygen glucose deprivation. Proceedings of the National Academy of Sciences 99, 15188–15193 (2002). 10.1073/pnas.232473399

11 Liang, R. et al. Discovery of Stable and Permeable N-Methylated Cyclic Peptides to Block the Endocytosis of GluA2 AMPAR for Ischemic Stroke Therapy. Journal of Medicinal Chemistry 69, 5594–5609 (2026). 10.1021/acs.jmedchem.5c02803

12 Collyer, E., Boyle, B. R., Gomez-Galvez, Y., Iacovitti, L. & Blanco-Suarez, E. Absence of chordin- like 1 aids motor recovery in a mouse model of stroke. Exp Neurol, 114548 (2023). 10.1016/j.expneurol.2023.114548

13 Koszegi, Z., Fiuza, M. & Hanley, J. G. Endocytosis and lysosomal degradation of GluA2/3 AMPARs in response to oxygen/glucose deprivation in hippocampal but not cortical neurons. Scientific Reports 7, 12318 (2017). 10.1038/s41598-017-12534-w

14 Blanco-Suarez, E., Liu, T. F., Kopelevich, A. & Allen, N. J. Astrocyte-Secreted Chordin-like 1 Drives Synapse Maturation and Limits Plasticity by Increasing Synaptic GluA2 AMPA Receptors. Neuron 100, 1116–1132.e1113 (2018). 10.1016/j.neuron.2018.09.043

15 Blanco-Suarez, E. & Allen, N. J. Astrocyte-secreted chordin-like 1 regulates spine density after ischemic injury. Sci Rep 12, 4176 (2022). 10.1038/s41598-022-08031-4

16 Blanco-Suárez, E. in Neural Repair: Methods and Protocols (eds Vardan T. Karamyan & Ann M. Stowe) 29–38 (Springer US, 2023).

17 Boyle, B. R., Collyer, E., Berghella, A. P. & Blanco-Suarez, E. Protocol for inducing focal striatal stroke in mice using photothrombosis. STAR Protoc 7, 104487 (2026). 10.1016/j.xpro.2026.104487

18 Schmidt-Kastner, R. Genomic approach to selective vulnerability of the hippocampus in brain ischemia-hypoxia. Neuroscience 309, 259–279 (2015). 10.1016/j.neuroscience.2015.08.034

19 Chan, K. Y. et al. Engineered AAVs for efficient noninvasive gene delivery to the central and peripheral nervous systems. Nature Neuroscience 20, 1172–1179 (2017). 10.1038/nn.4593

20 Challis, R. C. et al. Systemic AAV vectors for widespread and targeted gene delivery in rodents. Nat Protoc 14, 379–414 (2019). 10.1038/s41596-018-0097-3

21 Zemla, R. & Basu, J. Hippocampal function in rodents. Curr Opin Neurobiol 43, 187–197 (2017). 10.1016/j.conb.2017.04.005

22 Padilla-Coreano, N. et al. Direct Ventral Hippocampal-Prefrontal Input Is Required for Anxiety- Related Neural Activity and Behavior. Neuron 89, 857–866 (2016). 10.1016/j.neuron.2016.01.011

23 Schmidt, A. et al. Progressive cognitive deficits in a mouse model of recurrent photothrombotic stroke. Stroke 46, 1127–1131 (2015). 10.1161/strokeaha.115.008905

24 Blanco-Suárez, E., Fiuza, M., Liu, X., Chakkarapani, E. & Hanley, J. G. Differential Tiam1/Rac1 activation in hippocampal and cortical neurons mediates differential spine shrinkage in response to oxygen/glucose deprivation. Journal of Cerebral Blood Flow & Metabolism 34, 1898–1906 (2014). 10.1038/jcbfm.2014.158

25 Barth, A. M. & Mody, I. Changes in hippocampal neuronal activity during and after unilateral selective hippocampal ischemia in vivo. J Neurosci 31, 851–860 (2011). 10.1523/jneurosci.5080-10.2011

26 Arundine, M. & Tymianski, M. Molecular mechanisms of calcium-dependent neurodegeneration in excitotoxicity. Cell Calcium 34, 325–337 (2003). 10.1016/S0143-4160(03)00141-6

27 Choi, D. W. Excitotoxicity: Still Hammering the Ischemic Brain in 2020. Frontiers in Neuroscience 14, 1104 (2020).

28 Lai, T. W., Zhang, S. & Wang, Y. T. Excitotoxicity and stroke: Identifying novel targets for neuroprotection. Progress in Neurobiology 115, 157–188 (2014). 10.1016/j.pneurobio.2013.11.006

29 Wang, F., Xie, X., Xing, X. & Sun, X. Excitatory Synaptic Transmission in Ischemic Stroke: A New Outlet for Classical Neuroprotective Strategies. Int J Mol Sci 23 (2022). 10.3390/ijms23169381

30 Dobkin, B. H. & Carmichael, S. T. The Specific Requirements of Neural Repair Trials for Stroke. Neurorehabilitation and Neural Repair 30, 470–478 (2015). 10.1177/1545968315604400

31 Carmichael, S. T. Brain Excitability in Stroke: The Yin and Yang of Stroke Progression. Archives of Neurology 69, 161–167 (2012). 10.1001/archneurol.2011.1175

32 Lu, W. et al. Subunit composition of synaptic AMPA receptors revealed by a single-cell genetic approach. Neuron 62, 254–268 (2009). 10.1016/j.neuron.2009.02.027

33 Noh, K.-M. et al. Blockade of calcium-permeable AMPA receptors protects hippocampal neurons against global ischemia-induced death. Proceedings of the National Academy of Sciences of the United States of America 102, 12230–12235 (2005). 10.1073/pnas.0505408102

34 Gleichman, A. J., Kawaguchi, R., Sofroniew, M. V. & Carmichael, S. T. A toolbox of astrocyte- specific, serotype-independent adeno-associated viral vectors using microRNA targeting sequences. Nat Commun 14, 7426 (2023). 10.1038/s41467-023-42746-w

35 Farhy-Tselnicker, I. et al. Activity-dependent modulation of synapse-regulating genes in astrocytes. eLife 10, e70514 (2021). 10.7554/eLife.70514

36 Kraeuter, A. K., Guest, P. C. & Sarnyai, Z. The Open Field Test for Measuring Locomotor Activity and Anxiety-Like Behavior. Methods Mol Biol 1916, 99–103 (2019). 10.1007/978-1-4939-8994-2_9

37 Gawel, K., Gibula, E., Marszalek-Grabska, M., Filarowska, J. & Kotlinska, J. H. Assessment of spatial learning and memory in the Barnes maze task in rodents-methodological consideration. Naunyn Schmiedebergs Arch Pharmacol 392, 1–18 (2019). 10.1007/s00210-018-1589-y

38 Demichev, V., Messner, C. B., Vernardis, S. I., Lilley, K. S. & Ralser, M. DIA-NN: neural networks and interference correction enable deep proteome coverage in high throughput. Nat Methods 17, 41–44 (2020). 10.1038/s41592-019-0638-x

39 Di Tommaso, P. et al. Nextflow enables reproducible computational workflows. Nat Biotechnol 35, 316–319 (2017). 10.1038/nbt.3820

40 Fabian Pedregosa, G. V., Alexandre Gramfort, Vincent Michel, Bertrand Thirion, Olivier Grisel, Mathieu Blondel, Peter Prettenhofer, Ron Weiss, Vincent Dubourg, Jake Vanderplas, Alexandre Passos, David Cournapeau, Matthieu Brucher, Matthieu Perrot, Édouard Duchesnay. Scikit-learn: Machine Learning in Python. Journal of Machine Learning Research 12(85), 2825−2830 (2011).

41 Ritchie, M. E. et al. limma powers differential expression analyses for RNA-sequencing and microarray studies. Nucleic Acids Res 43, e47 (2015). 10.1093/nar/gkv007

42 Zhu, Y. et al. DEqMS: A Method for Accurate Variance Estimation in Differential Protein Expression Analysis. Mol Cell Proteomics 19, 1047–1057 (2020). 10.1074/mcp.TIR119.001646

43 Fang, Z., Liu, X. & Peltz, G. GSEApy: a comprehensive package for performing gene set enrichment analysis in Python. Bioinformatics 39 (2023). 10.1093/bioinformatics/btac757

